# The Arabidopsis Framework Model version 2 predicts the organism-level effects of circadian clock gene mis-regulation

**DOI:** 10.1101/105437

**Authors:** Yin Hoon Chew, Daniel D. Seaton, Virginie Mengin, Anna Flis, Sam T. Mugford, Gavin M. George, Michael Moulin, Alastair Hume, Samuel C. Zeeman, Teresa B. Fitzpatrick, Alison M. Smith, Mark Stitt, Andrew J. Millar

## Abstract

Predicting a multicellular organism’s phenotype quantitatively from its genotype is challenging, as genetic effects must propagate across scales. Circadian clocks are intracellular regulators that control temporal gene expression patterns and hence metabolism, physiology and behaviour. Here we explain and predict canonical phenotypes of circadian timing in a multicellular, model organism. We used diverse metabolic and physiological data to combine and extend mathematical models of rhythmic gene expression, photoperiod-dependent flowering, elongation growth and starch metabolism within a Framework Model for the vegetative growth of *Arabidopsis thaliana*, sharing the model and data files in a structured, public resource. The calibrated model predicted the effect of altered circadian timing upon each particular phenotype in clock-mutant plants under standard laboratory conditions. Altered night-time metabolism of stored starch accounted for most of the decrease in whole-plant biomass, as previously proposed. Mobilisation of a secondary store of malate and fumarate was also mis-regulated, accounting for any remaining biomass defect. We test three candidate mechanisms for the accumulation of these organic acids. Our results link genotype through specific processes to higher-level phenotypes, formalising our understanding of a subtle, pleiotropic syndrome at the whole-organism level, and validating the systems approach to understand complex traits starting from intracellular circuits.

This work updates the first biorXiv version, February 2017, https://doi.org/10.1101/105437, with an expanded description and additional analysis of the same core data sets and the same FMv2 model, summary tables and supporting, follow-on data from three further studies with further collaborators. This biorXiv revision constitutes the second version of this report.

## Introduction

Circadian clocks in all organisms integrate multiple environmental inputs and affect disparate, potentially interacting biological processes, from sleep/wake cycles in mammals to flowering in plants (Kuhlman *et al.* 2018). Clock genes are rarely essential but appropriate alleles can confer an organismal growth (Green *et al.* 2002) and competitive advantage (Dodd *et al.* 2005, Ouyang *et al.* 1998) and have been repeatedly selected during crop domestication (Bendix *et al.* 2015, Muller *et al.* 2015). Mistimed mutant organisms suffer a syndrome of mild, environment-dependent effects akin to a chronic disease (Dodd *et al.* 2005, Paschos *et al.* 2010, Peek *et al.* 2013), including traits that are not overtly related to rhythmicity. Small networks of “clock genes” drive these 24-hour, biological rhythms in eukaryotes (Kuhlman *et al.* 2018, Millar 2016). A few among thousands of downstream, clock-regulated genes are known to mediate physiological phenotypes, such as the metabolic syndrome of clock mutant animals (Peek *et al.* 2013). However, identifying such causal links along biosynthetic, signalling or developmental pathways is not sufficient to predict quantitatively the whole-organism phenotypes due to circadian mis-timing: formal, mathematical models are required.

Predictive modelling from cellular mechanisms in multicellular organisms has best succeeded for phenotypes that closely map the behaviour of gene circuits (von Dassow *et al.* 2000) or signalling pathways (Band *et al.* 2014). From the whole-organism perspective, these cases can be described as having a “short phenotypic distance” from the genotype (Hammer *et al.* 2019). However, a recent review (Clark *et al.* 2020) points out that whole-plant models that link to the molecular level “are rare (Chew *et al.* 2014, Grafahrend-Belau *et al.* 2013)”, despite the long-standing aspiration to understand and manipulate the field-scale traits that distinguish crop varieties based on their underlying genetic differences (Hammer *et al.* 2019, Matthews and Marshall-Colon 2021), and the potential applications in breeding programmes or studies of natural selection. Here, we extend one of the rare, whole-plant models to understand the pleiotropic phenotypes controlled by the circadian clock.

The Arabidopsis Framework Model version 1 (FMv1) represents the interacting, physiological components of vegetative growth in *Arabidopsis thaliana* up to flowering, in a simple, modular fashion (Chew *et al.* 2014). The model was designed to study circadian effects on physiology but has proved more broadly useful (Matthews and Marshall-Colon 2021), as evidenced by the re-use and adaptation of FMv1 to understand the effects of gibberellin-related regulatory proteins on biomass and leaf architecture (Pullen *et al.* 2019), of temperature on flowering *via FT* gene regulation and/or leaf development (in FMv1.5, Kinmonth-Schultz *et al.* 2019), and of phytochrome-regulated cotyledon size upon adult plant biomass (Krahmer *et al.* 2021). A simplified version of FMv1 (FM-lite) was included in a whole-lifecycle model of Arabidopsis (FM-life) to test the adaptive value of phenology traits (Zardilis *et al.* 2019). We now update FMv1 with biochemically-detailed submodels of the circadian clock and of its outputs, and a revised allocation of photosynthetic carbon with a circadian input.

The Arabidopsis clock mechanism comprises dawn-expressed transcription factors *LATE ELONGATED HYPOCOTYL* (*LHY*) and *CIRCADIAN CLOCK-ASSOCIATED 1* (*CCA1*), which inhibit the expression of evening genes such as *GIGANTEA* (*GI*), as illustrated by their respective RNA expression profiles in plants grown under light:dark cycles (Figure 1a) (Flis *et al.* 2015). *LHY* and *CCA1* expression is inhibited by PSEUDO-RESPONSE REGULATOR (PRR) proteins. Removing the earliest-expressed *PRR* genes in *prr7prr9* mutants slows the clock (Nakamichi *et al.* 2010), because *LHY* and *CCA1* expression is not inhibited as quickly as in wild-type plants. Under constant light, rhythmic outputs have a circadian period around 30h in the double-mutant plants, compared to 24.5h in the wild type (Salome and McClung 2005). The decline of *LHY* and *CCA1* RNA levels in the double mutants is delayed under light:dark cycles, as is the subsequent rise of *GI* and other target genes (Figure 1b). A series of mathematical models has represented the increasingly detailed biochemistry of the clock gene circuit (Fogelmark and Troein 2014, Pokhilko *et al.* 2012, Urquiza-García and Millar 2021), recapitulating the effects of the *prr7prr9* mutations on clock gene dynamics (Figs. 1a,1b) (Flis *et al.* 2015, Pokhilko *et al.* 2012).

**Figure 1:**
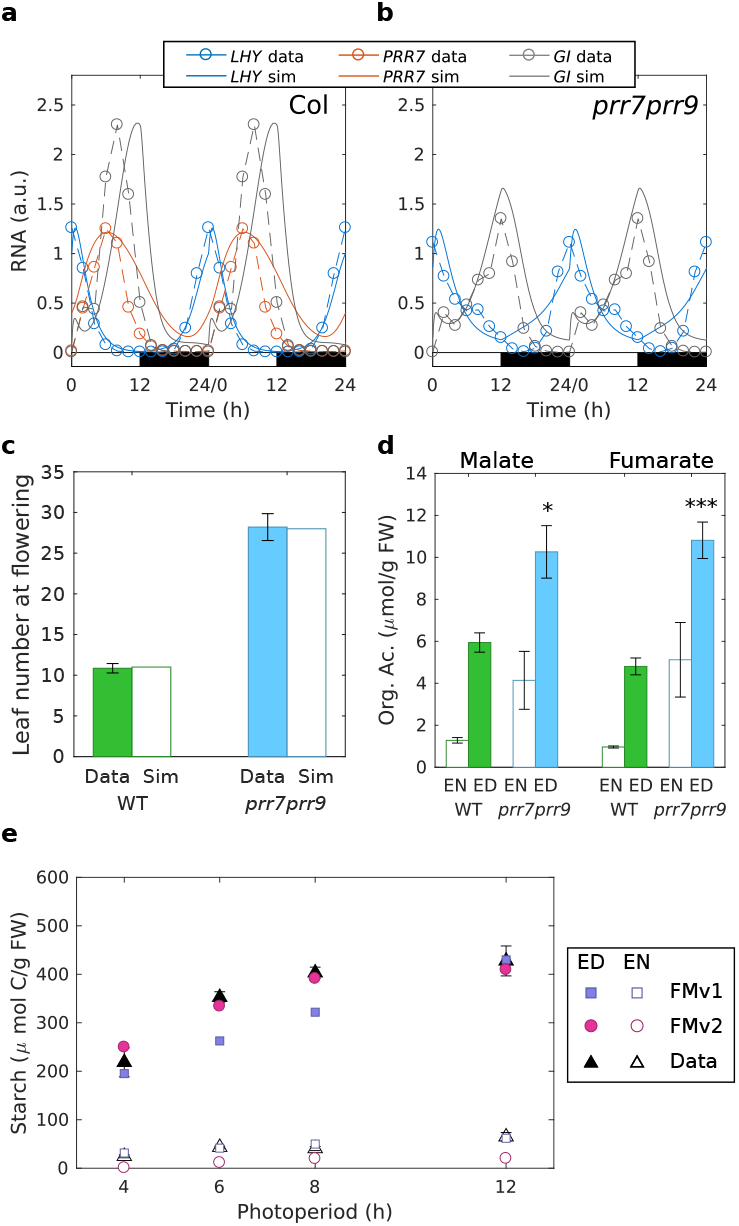
Simulation of clock dynamics and clock outputs. (a,b) Clock gene mRNA abundance (Flis *et al.* 2015) for wild-type (Col) and *prr7prr9* plants (dashed lines, symbols), and FMv2 simulations (solid lines), under 12h light: 12h dark cycles (12L:12D), double-plotted, normalised to Col level. (c) Rosette leaf number at flowering(Nakamichi *et al.* 2007) under 16L:8D (filled), compared to simulation (open). (d) Malate and fumarate accumulation (mean±SEM, n=4) in Col and *prr7prr9* at end of day (ED) or night (EN) under 12L:12D, 20°C, light intensity=160 *μ*mol/m^2^/s; t-tests compared *prr7prr9* to Col (* p<0.05; *** p<0.001). (e) Starch levels at ED (filled) and EN (open) after 30 days under various photoperiods(Sulpice *et al.* 2014) (triangles), compared to FMv1 (squares), FMv2 (circles).

The clock models have been extended to the output pathways that convey circadian timing information to control downstream processes, starting with the control of flowering time in response to photoperiod (Salazar *et al.* 2009, Welch *et al.* 2005). The light-responsive pathways of gene regulation that control both flowering and organ elongation have been modelled with increasing biochemical detail (Seaton *et al.* 2015), while other models have represented the non-circadian inputs that also control flowering (Chew *et al.* 2012, Wilczek *et al.* 2009). The first representations of circadian metabolic outputs have focussed on the nightly, clock-limited rate of sugar mobilisation from storage in transient starch (Graf *et al.* 2010). Starch receives the largest portion of photosynthate in Arabidopsis leaves, after sugars, and clock regulation ensures that leaves use as much of each day’s carbon store for night-time growth as possible, leaving just 2-5% at dawn (Smith and Zeeman 2020). Models of this process have focussed variously on the biochemistry of the regulated degradation, its connection to the clock mechanism, or the potential impact of the resulting sugar on clock dynamics (Pokhilko *et al.* 2014, Scialdone *et al.* 2013, Seaton *et al.* 2014, Seki *et al.* 2017), but without considering the resulting growth of the plant. We therefore tested whether these proposed mechanisms were sufficient to understand the multiple phenotypes of the long-period clock mutant *prr7prr9*, not only for the photoperiodic regulation of flowering, which is arguably a short phenotypic distance from the clock genes, but also the more distant biomass growth and metabolic traits.

## Results

### Circadian control of plant development

The FMv1 was designed with a modular structure, which facilitated our replacement of the earlier clock and photoperiod pathway submodel (Salazar *et al.* 2009) with more detailed submodels (Pokhilko *et al.* 2012, Seaton *et al.* 2015) that explicitly represent the clock genes *PRR7*, *PRR9* and output pathway genes (see Supporting Information, section 1). Table 1 summarises the changes in the submodels of the Arabidopsis Framework Model version 2 (FMv2) compared to FMv1, and notes the abbreviated names for the clock (P2011), starch (S2014) and photoperiodism (S2015) submodels. Figure 2 shows the updated gene circuits in the context of the Framework Model’s other functions.

**Table 1.** Changes to the submodels and processes of the FMv2 compared to the FMv1. The origins of the multiple submodels that were combined to form the FMv1 are noted, together with the origins of the updated submodels and other components of the FMv2.

| Model function | Process | In FMv1 | Derived from | In FMv2 | Sub-model | Derived from |
| --- | --- | --- | --- | --- | --- | --- |
| Environmental data input | Hourly Light intensity, [CO<sub>2</sub>], temperature, sunrise/sunset times |  | - | as in FMv1 |  |  |
| Functional structural plant model | Organ sink demand | Thermal time function | (Christophe *et al.* 2008) | as in FMv1 |  |  |
|  | Leaf initiation | Stochastic function | - | as in FMv1 |  |  |
|  | Shoot/root allocation | Emergent behaviour | - | as in FMv1 |  |  |
| Carbon dynamic model | Photosynthate partitioning | "Basal + Overflow" | (Rasse and Tocquin 2006) | "Starch-first", using AGPase data | - | this work |
|  | Starch mobilisation | Fixed rate | (Rasse and Tocquin 2006) | Controlled by P2011 clock model | from S2014 Model 2 | (Seaton *et al.* 2014) |
|  | Malate and Fumarate pool | - | - | New model component | - | this work |
| Circadian clock model | Light-entrained rhythms | 4 RNAs, 4 proteins, 0 protein complexes; lacks *PRR7*, *PRR9* | (Locke *et al.* 2005) | 9 RNAs, 16 proteins, 6 protein complexes; includes *PRR7*, *PRR9* | P2011 | (Pokhilko *et al.* 2012) |
| Photoperiodism model | External coincidence detector | 1 RNA, 1 Protein | (Salazar *et al.* 2009) | 4 RNAs, 3 Proteins | S2015 | (Seaton *et al.* 2015) |
| Light signalling model | Phytochrome/PIF pathway | - | - | 4 RNAs, 3 Proteins | S2015 | (Seaton *et al.* 2015) |
| Phenology model | Seedling emergence, flowering time | Diel photothermal time | (Chew *et al.* 2012) | as in FMv1 |  |  |

**Figure 2.**
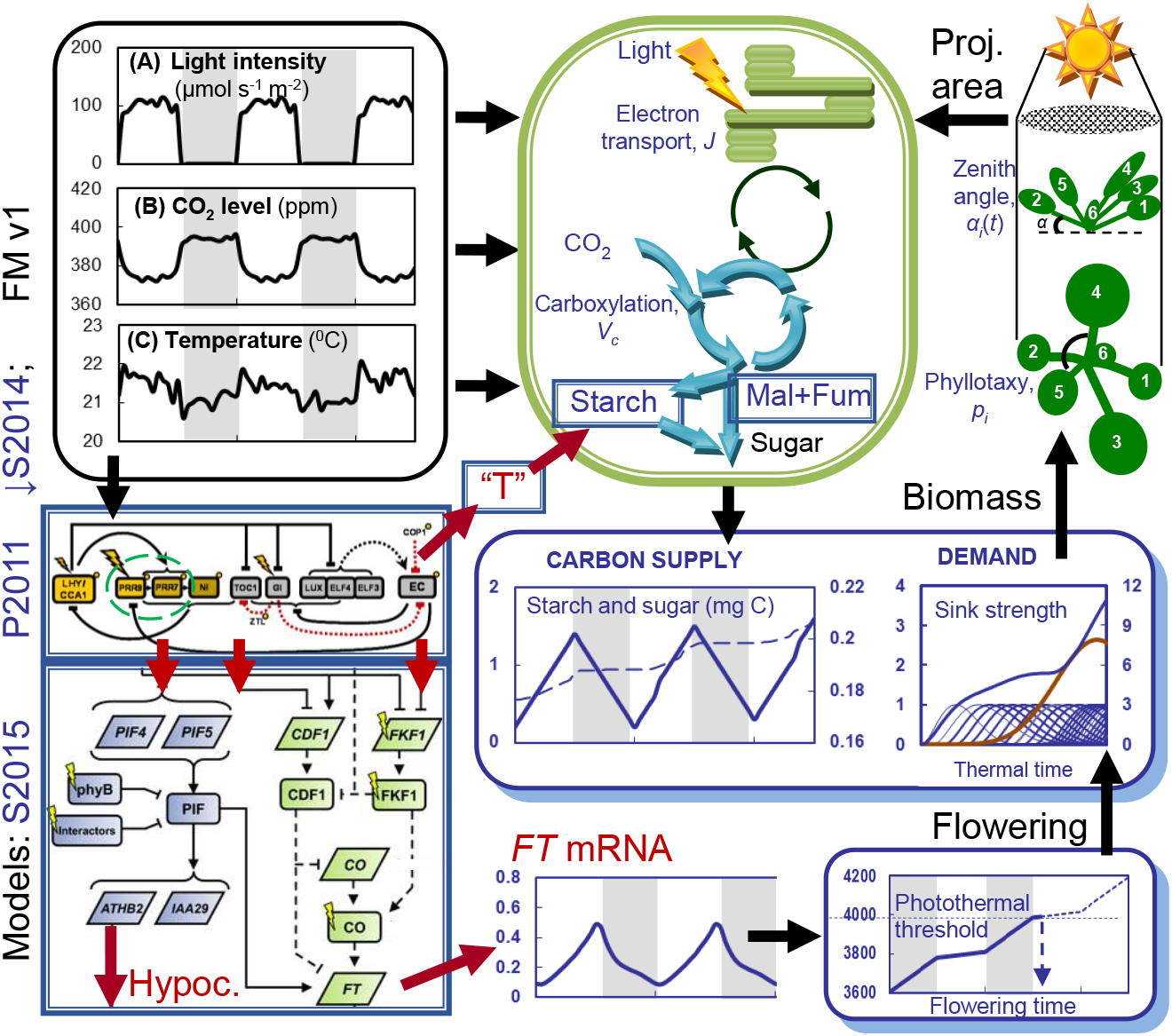
Operation of the Arabidopsis Framework Model version 2 (FMv2). Model components that are updated in FMv2 are bounded in blue rectangles, with abbreviated model names (left). Key outputs from the new models are shown as red arrows. The P2011 clock model shows only clock genes for clarity. Morning or day genes, yellow symbols; evening genes and the Evening Complex (EC), grey symbols. The PRR9 and PRR7 genes that are inactivated in the prr7prr9 double mutant are marked with a dashed green oval. Components of P2011 drive the S2014 starch degradation model via component “T” (centre), and the S2015 external coincidence model (lower left). S2015 controls photoperiod-dependent hypocotyl elongation (Hypoc.) by rhythmically gating the PhyB/PIF/ATHB2 pathway, and the florigen FT by the CDF1/FKF1/CO pathway. The S2015 cartoon distinguishes RNA components (paralellograms) from proteins (rounded rectangles). Light inputs are shown as flashes. Among the model components retained from the FMv1, the Carbon Dynamic Model (bounded in green) includes updated starch and malate and fumarate (Mal+Fum) carbon stores, and provides Sugar as the Carbon Supply for growth. This is allocated according to the Demand of leaf (blue) and root (red) sinks in the Functional-Structural Plant Model, which uses the leaf Biomass to calculate the rosette’s projected area for photosynthesis (Proj. area). When the Photothermal model (lower right) reaches the threshold for Flowering, the simulation ends.

Flowering time in Arabidopsis is commonly scored by the number of rosette leaves produced before the flowering bolt elongates above the vegetative rosette, under long and short photoperiods. Predicting leaf number involves the FM’s clock and photoperiod (Seaton *et al.* 2015), phenology (Chew *et al.* 2012) and functional-structural sub-models (Christophe *et al.* 2008), suggesting that this phenotype can only be considered a “short distance” from the clock genes in the sense that it doesn’t involve the metabolic submodel. The P2011 clock sub-model (Pokhilko *et al.* 2012) was known to recapitulate the altered clock gene expression dynamics in *prr7prr9* double mutant plants in general. Figures 1a, 1b show that simulations of the published model also closely matched the later, reference dataset of RNA timeseries from the TiMet project (Flis *et al.* 2015) in the double mutants. In brief, the late circadian phase in the mutant simulation reduces the *FT* mRNA output from the S2015 submodel (see Supporting Information, section 1; Figure 2). The phenology model therefore takes longer to reach the abstract photothermal threshold for flowering, giving more time for the functional-structural model to produce rosette leaves.

The new model combination in the FMv2 matched the leaf number data published by the Mizuno group (Nakamichi *et al.* 2007) for wild-type Columbia (Col) plants under long-photoperiod growth conditions (Figure 1c). The expectation from FMv1 was that calibration of the photothermal time threshold for flowering (Figure 2) might often be required for the model to match the absolute leaf numbers observed across different laboratories (Chew *et al.* 2014) but that calibration was not required in this case. The late circadian phase predicted a lower activation of the photoperiod response mechanism and predicted a late-flowering phenotype in the *prr7prr9* mutants in these conditions, which was also exactly matched by the model (Figure 1c). Under short photoperiods, wild-type plants flower late and the clock-dependent, photoperiodic pathway has little effect. It is therefore expected that the double mutant’s flowering time is closer to the wild type, as observed in the data and qualitatively in the simulations (Supporting Information Figure 1a). The model predicted slightly later flowering than was observed in both genotypes, however, suggestive of a non-photoperiodic effect on flowering.

The S2015 submodel further predicts the regulation of elongation growth in the seedling hypocotyl, which is light-and therefore photoperiod-dependent. The FMv2 matched the observed photoperiodic regulation of hypocotyl elongation in wild-type plants, in data again from the Mizuno group (Niwa *et al.* 2009). The longer hypocotyls of *prr7prr9* seedlings were qualitatively matched by the model (Supporting Information Figure 1b), with an altered balance between long and short photoperiods that suggests further, secondary mechanism(s) may be involved.

### Mis-regulation of carbon dynamics

In contrast to the rapid, finite elongation of the seedling hypocotyl, long-term biomass growth of the whole plant is mediated by the metabolic network, the development of sink and source organs and the partitioning of metabolic resources amongst them. Among many potential circadian effects on metabolism, we focus on the clock-limited rate of sugar mobilisation from starch. First, we addressed an observed limitation of the starch dynamics in the metabolic sub-model of the FMv1. Daytime starch accumulation in wild-type plants was underestimated by the model under short photoperiods (Chew *et al.* 2014, Sulpice *et al.* 2014), indicating that the ‘growth-first’ control of photoassimilate partitioning in FMv1 should be updated. Instead of an ‘overflow’ to starch after growth demand was satisfied, FMv2 uses photoperiod-dependent, ‘starch-first’ partitioning. The key, starch biosynthetic enzyme, AGPase, was clearly photoperiod-dependent at the level of measured activity (Supporting Information Figure 2a)(Mugford *et al.* 2014), and in the abundance of some protein subunits (Seaton *et al.* 2018). The measured photoperiod control of AGPase activity was therefore introduced into the model as a candidate mechanism for this photoperiod-dependent partitioning (Supporting Information, section 2). This change partitioned sufficient carbon to starch in simulated, short photoperiods to match the experimental data in photoperiods of up to 12h (Supporting Information Figure 2a). Under longer photoperiods, Arabidopsis plants manage the additional photosynthate using additional metabolic regulation (Sulpice *et al.* 2014) that is not represented in the FMv2.

At night, starch is mobilised (degraded) at a constant rate to provide sugar until dawn, as anticipated by the circadian clock (Graf *et al.* 2010, Scialdone *et al.* 2013). We therefore included the starch degradation mechanism from a further, published sub-model (S2014) that links starch degradation to the clock sub-model, and had been shown to match the early depletion of starch reserves in mutant plants with a short circadian period and in other, discriminating conditions (Seaton *et al.* 2014) (Supporting Information, section 1). Simulation of the revised model closely matched the starch levels of wild-type plants under photoperiods of up to 12h (Figure 1e). Finally, the organic acids malate and fumarate also accumulate significantly during the day in Arabidopsis, are mobilised at night and have been proposed as secondary carbon stores (Zell *et al.* 2010). Analysis of 22 further metabolites revealed that at the end of the day, levels of malate and fumarate were two-fold higher in *prr7prr9* mutants than wild type, with a lesser elevation in citrate and in the much smaller pools of aconitate and iso-citrate (Figure 1d, Supporting Information Figure 3). The malate and fumarate pools were therefore prioritised for further analysis in the double mutants, alongside starch. As a simple approximation, the sum of malate and fumarate was included as an “organic acid” pool with dynamics similar to starch, in the FMv2 (see Supporting Information, section 3; Figure 2). If altered starch mobilisation in the long-period clock mutant *prr7prr9* was sufficient to affect its biomass, then the FMv2 should also predict that phenotype.

### Control of biomass via starch degradation

We first tested whether the FMv2 could explain the biomass phenotype caused by a mild, direct change in starch degradation, in mutants of *LIKE SEX FOUR 1 (LSF1)* that were tested for 12 major metabolite pools (Arrivault *et al.* 2009, Pyl *et al.* 2012) and the physiological parameters required to calibrate the FMv2. *LSF1* encodes a phosphatase homologue located in the chloroplast, which interacts with beta-amylases and is necessary for normal starch mobilisation (Comparot-Moss *et al.* 2010, Schreier *et al.* 2019). *lsf1* mutants grown under 12L:12D have mildly elevated starch levels and reduced biomass, similar to the *prr7prr9* clock mutant (Figs.3b,3c,3e). Reducing the relative starch degradation rate in the FMv2 (Supporting Information Table 1) recapitulated the *lsf1* starch excess observed in published studies (Comparot-Moss *et al.* 2010) (Supporting Information Figure 1c) and in new data (Fig.3e). To reproduce the biomass growth of wild-type plants, a minimal model calibration workflow (Supporting Information Figure 4) used measured photosynthetic and metabolic variables (Supporting Information Figure 5) to calibrate the model for our conditions (Supporting Information Table 1). The water content was previously highlighted as a key parameter; by setting its value to the mean water content observed for each genotype, model predictions can be compared to fresh weight data (Chew *et al.* 2014). Introducing the lower starch degradation observed in *lsf1* and its measured water content into the calibrated model accurately predicted the reduced biomass of *lsf1* mutant plants (Fig.3c). The coefficient of variation of the Root-Mean-Square Error (cvRMSE) provides a normalised error metric for all biomass data (Chew *et al.* 2014), showing a good fit to both *lsf1* and wild-type genotypes (8.0% Col and 13.7% *lsf1)*, and validating the FMv2 for biomass prediction. In addition to the direct prediction of carbon biomass, the FMv2 predicts the gain of other major biomass components indirectly, allowing further validation. The change in biomass gain from day to night in simulated, wild-type plants predicted a 3.3-fold increase in protein synthesis rates in daytime, compared to night-time, for example. This was very close to the observed 3.1-fold increase in protein synthesis in published data (Ishihara *et al.* 2015)(see Supporting Information, section 5).

**Figure 3:**
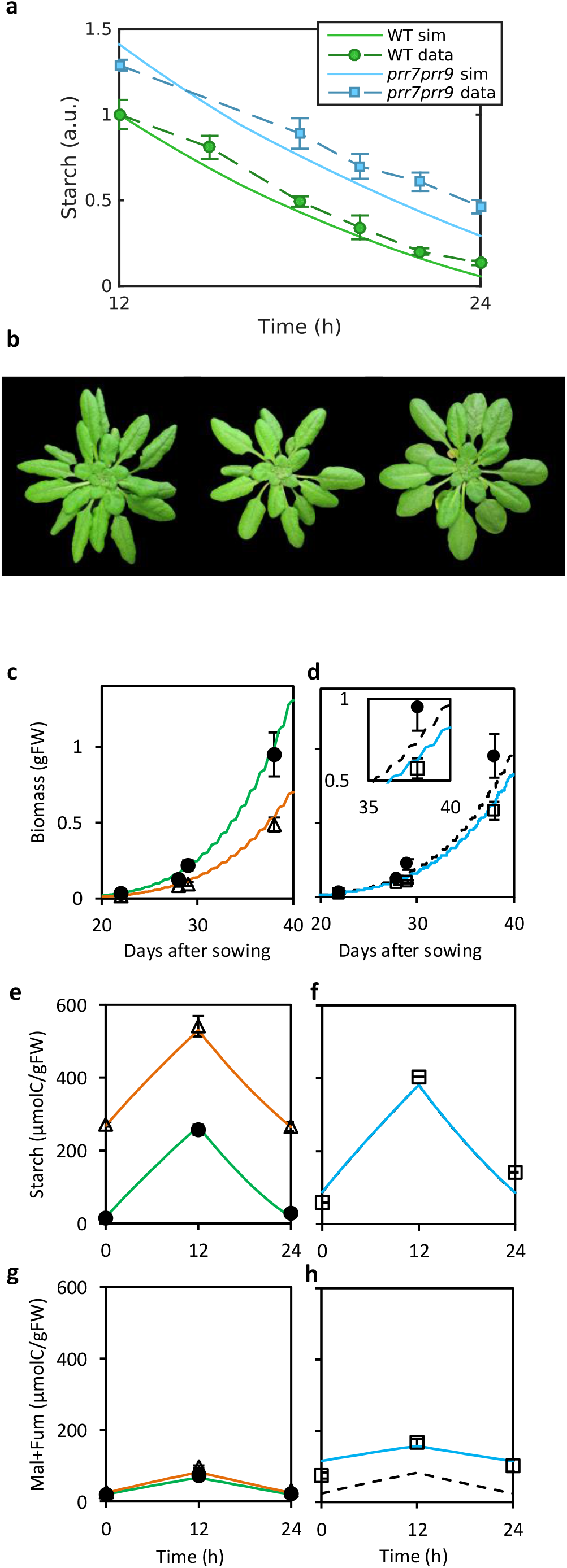
Contributions of starch and organic acids to biomass growth. (a) *prr7prr9* plants (blue, squares) mobilised starch more slowly than Col (WT, green, circles), mean±SEM, n=6, giving a starch excess similar to simulations of the FMv2 (solid lines). Data and model normalised to Col at 12h. (b) 38-day-old Col, *lsf1, prr7prr9* plants. (c-h) Data (symbols) and simulation (lines) of fresh weight (c-d), starch (e-f) and total malate and fumarate (g-h) for Col (circles, green), *lsf1* (triangles, orange) and *prr7prr9* plants (squares). Simulations of *prr7prr9* show a starch defect only (dashed black line) or both starch and malate+fumarate defects (solid blue). (d) Inset enlarges main panel around 38-day data point. Data show mean±SD; n=5 for biomass; n=3 for metabolites, where each sample pooled 3 plants. Growth conditions, 12L:12D with light intensity=190 *μ*mol/m^2^/s (a) or 145 *μ*mol/m^2^/s (b-h), temperature 20°C (a), 20.5°C (b-h); CO_2_=420 ppm.

### Control of biomass in the circadian clock mutant

*prr7prr9* mutants showed slower relative starch degradation (Figure 3a) and higher starch levels at both dawn and dusk (Supporting Information Figure 1d) than the wild type. The starch profiles of plants grown in Norwich (Figure 3a, Supporting Information Figure 1d) and in Edinburgh (Figure 3f) closely matched the profiles predicted by simulating *prr7prr9* mutations in the clock sub-model of the FMv2. The biomass of *prr7prr9* mutant plants was 40% lower than wild-type control plants at 38 days. The calibrated FMv2 also predicted a lower biomass in *prr7prr9* due to the starch defect but with a smaller effect than in the data (26% biomass reduction). A poor model fit (cvRMSE = 41%) indicated that process(es) additional to starch degradation were likely to account for the limited growth of *prr7prr9* but not of *lsf1* plants in this study. *prr7* single mutants fully mobilised starch and grew normally, as predicted from their normal circadian timing (Supporting Information Figure 6). The mild starch excess at the end of the day aligns with past reports (Haydon *et al.* 2013, Seki *et al.* 2017) but apparently had no effect on biomass growth in *prr7* plants. We therefore sought another clock-regulated process that might contribute to reduce the biomass of *prr7prr9* plants.

Model calibration data showed that photosynthesis, starch synthesis and leaf production rates were unaffected by the *prr7prr9* mutations (Supporting Information Figure 5). Water content was slightly reduced in *prr7prr9* (Supporting Information Table 1) and this is the most sensitive parameter in our model (Supporting Information Figure 7). However, neither 1 S.D. variation in the mutant’s simulated water content, nor any measured water content value allowed the model with only the clock (and hence starch) defect to match the mutant biomass (Supporting Information Figure 8).

Considering the pool of malate and fumarate as a secondary carbon store (Zell *et al.* 2010), the amount of carbon mobilised from malate and fumarate at night in the wild type was up to 19% of the carbon mobilised from starch. *prr7prr9* but not *lsf1* plants accumulated excess malate and fumarate, representing further ‘wasted’ carbon that did not contribute to biomass growth (Figures 3g, 3h), consistent with independent sampling of this double mutant (Flis *et al.* 2019). We therefore reduced the relative malate and fumarate mobilisation rate in the FMv2 simulation of *prr7prr9*, to reproduce the observed organic acid excess (Figure 3h). Together, the simulated defects in starch and organic acid mobilisation quantitatively accounted for the mutant’s reduced biomass (Figure 3d; cvRMSE = 15.1%).

### Replication of the physiological phenotypes

The *prr7prr9* double mutants showed the phenotypes noted above in three further experiments (Figure 4). The final biomass of the double mutants ranged from 53% to 76% of the wild-type value, compared to a mean of 60% in experiment 1 (Figure 3). Calibrating the models with measured physiological parameters facilitated the comparison among these studies (see Supporting Information, section 4). The biomass of the double mutants in experiment 2 was also below the biomass predicted from slower starch degradation rate alone (Supporting Information Figure 9g). As in experiment 1, the photoassimilate retained in the mutant’s starch, malate and fumarate pools together accounted for its lower biomass in experiment 2. Both wild-type and clock-mutant genotypes grew slightly larger in experiment 3 than the model simulations predicted (Supporting Information Figure 9a). Photosynthetic rates measured at a single timepoint might not have adequately calibrated the model in this case to simulate growth over several weeks. The mutant’s starch degradation rate matched the FMv2 prediction in experiment 3 and the consequent, clock-dependent effect on starch degradation alone in the simulations sufficiently accounted for the mutants’ biomass (72% of wild-type biomass, cvRMSE 4.4%). Reducing the simulated mobilisation of the malate and fumarate pool, as in the simulations of experiments 1 and 2, slightly underestimated the mutants’ biomass in experiment 3 (Supporting Information Figure 9a; cvRMSE 12.1%). The Col and *prr7prr9* control plants in an experiment on gibberellin effects (see below) again replicated these phenotypes (“no GA”, Figure 4, Supporting Information Figure 9). The GA study was not designed to collect full calibration data, so simulations were not performed.

**Figure 4:**
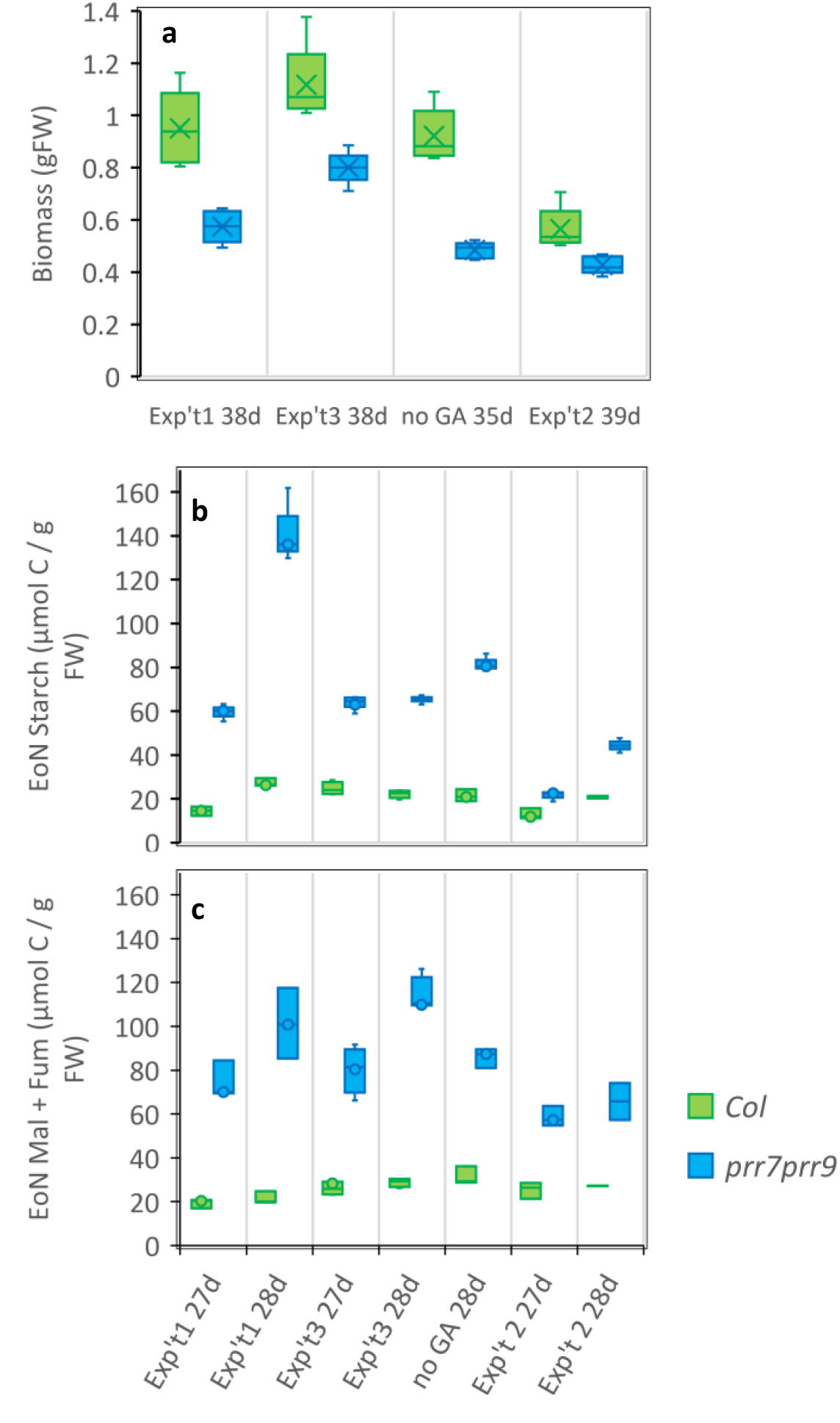
Replication of biomass and carbon storage traits. a) The range of *prr7prr9* final biomass (indicated by whiskers; blue) is below the range for Col (green) in all experiments, with the smallest difference in experiment 2. Data are mean, n=5, error bar = SD. Exp’t 1 and 3, 38 days after sowing (38d); no GA controls, 35 days; Exp’t 2, 39 days. *prr7prr9* plants also show excess levels of (b) starch and (c) malate + fumarate at the end of the night (EoN) in all experiments, with the starch level of *prr7prr9* plants closest to Col in experiment 2. Data are mean, n=3 (exp’t 3, n=4), error bar = SD (Exp’t 1, 2 and 3, 27 and 28 days after sowing; no GA controls, 28 days only). Data collated from Figure 3 and Supporting Information Figures 9 and 11. Differences in sampling among experiments are outlined in Supporting Information.

Experiment 2 showed slower biomass growth, and the smallest effects of the *prr7prr9* mutation on several phenotypes (Figure 4; Supporting Information Figure 9). Mutant plants retained mean starch levels over 60 μmolC/gFW at the end of the night in all studies except experiment 2, compared to a wild-type level of 14-27 μmolC/gFW (Figure 4b). In-chamber recordings showed aberrant temperature averaging 18.5°C in the growth chamber for this experiment alone, rather than the intended 20.5°C. The double-mutant clock’s long circadian period is known to be fully restored to wild type at 12.5°C (Salome *et al.* 2006), hence the lower growth temperature likely weakened the effect of this temperature-conditional mutation, in this experiment. The P2011 clock model is calibrated for regulation at 21°C and is not expected to simulate temperature-dependent effects, hence the modelling of experiment 2 used observed rather than predicted starch degradation rates (see Supporting Information, section 4; Supporting Information Table 1).

These results provide a proof of concept, showing that the pervasive effects of altered circadian timing at the level of the whole plant can be understood quantitatively, in terms of the underlying biochemical and molecular mechanisms (see Discussion). The effect of clock mutations on starch metabolism was most significant in accounting for the mutants’ lower biomass, and changes in malate and fumarate levels accounted for any remaining growth defect.

### Testing potential mechanisms for circadian misregulation of organic acid pools

We next tested several processes that might alter malate and fumarate synthesis or utilisation in *prr7prr9*. The FMv2 simulations reduced malate and fumarate utilisation in the *prr7prr9* mutant but we also tested the possibility of increased malate synthesis, *via* CO_2_ fixation by PEP carboxylase. This can be measured as ^14^CO_2_ fixation in darkness, which was very low compared with photosynthetic CO_2_ assimilation (Kölling *et al.* 2015) but nevertheless measurable. Total dark CO_2_ fixation was similar in the *prr7prr9* mutants and the wild type (Supporting Information Figure 10a). However, the distribution of labelled carbon between compound classes differed (Supporting Information Figures 10b, 10c). Most was found in water-soluble acidic compounds (i.e. organic acids derived from PEP carboxylase action) and water-soluble basic compounds (presumably amino acids produced from the organic acids). While *prr7prr9* had slightly less label in soluble compounds overall, there was more in the acidic fraction and less in the basic fraction than in the wild type. Some label was found in proteins, with the amount in *prr7prr9* (12 ± 1%) significantly exceeding that of the wild type (7% ± 1%) leading to a slightly higher labelling of the insoluble fraction in the mutant.

Second, altered expression of the clock-regulated, thiamine-biosynthetic gene *THIAMINE C (THIC)* in *prr7prr9* might alter organic acid levels, by affecting thiamine-cofactor-dependent, metabolic enzymes (Bocobza *et al.* 2013, Raschke *et al.* 2007, Rosado-Souza *et al.* 2019). Biosynthetic intermediates TMP and thiamine indeed accumulated at 2- to 3-fold higher levels in the *prr7prr9* mutants (Supporting Information Figure 10f, 10g), though levels of THIC protein were reduced in this genotype (Supplementary Table 2 of (Graf *et al.* 2017). The level of the active TDP cofactor (vitamin B1) increased by less than 10% (Supporting Information Figure 10e), which was not statistically significant and suggested that homeostatic mechanisms were largely compensating for any circadian mis-regulation.

Third, reduced carbon demand from growth, rather than or in addition to redirected carbon supply, might promote organic acid accumulation indirectly (see Discussion). We applied exogenous gibberellins (GAs) to test whether stimulating growth could increase biomass in *prr7prr9*, by mobilising the mutant’s excess carbon stores into biomass. GA treatment indeed increased the biomass of both genotypes to a similar extent (Supporting Information Figure 11a) but this was not through greater mobilisation of the clock-dependent carbon pools. GA treatment had little or no effect on starch or malate levels at the end of the night (Ribeiro *et al.* 2012) (Supporting Information Figure 11c-11e), and increased the fumarate level. The starch level at the end of the day was slightly reduced in GA-treated *prr7prr9* plants. To control for a direct effect of GA on the mutants’ clock defect, given that the clock affects GA sensitivity (Arana *et al.* 2011), we confirmed that GA treatment did not rescue the circadian period defect in *prr7prr9* plants tested by luciferase reporter gene imaging under constant light (Supporting Information Figure 11f). Thus GA treatment likely stimulated biomass growth by a mechanism orthogonal to the growth-limiting effect of the clock mutations. Consistent with this, we found little overlap in previously-tested transcriptome responses to GA treatment (Bai *et al.* 2012) and the *prr5prr7prr9* clock mutation (Nakamichi *et al.* 2009a) (Supporting Information Figure 11g).

## Discussion

The Arabidopsis Framework Model version 2 (FMv2) builds upon the delayed gene expression patterns in *prr7prr9* double mutants to predict canonical clock phenotypes, under the standard laboratory conditions previously used to define these phenotypes. The greater hypocotyl elongation and delayed flowering time simulated in the mutants result from gene expression cascades in the S2015 sub-model, which represents similar molecular processes to the clock gene circuit represented in the P2011 sub-model. This aspect of the FMv2 is helpful to understand the effects of these gene expression regulators at the whole-plant scale (Krahmer *et al.* 2019). Regulatory pathways beyond the light- and clock-gene network could be included in future, but the earlier FMv1 model has been most used to simulate the growth of the Arabidopsis rosette, or the interaction of rosette development with these gene circuits. The FMv2 bridges from gene circuit dynamics not only to these developmental issues but also to carbon biomass growth, *via* its simple model of photosynthetic metabolism. Supporting Information Table 2 summarises the predictions tested here, the gaps revealed and those that could be tested in future.

Rhythmic output from the clock gene circuit to the metabolic network, in the chloroplast, controls the simulated level of transient starch that remains at the end of the night in the FMv2. The model correctly predicted the slower relative starch degradation rate, due to the delayed circadian timing of the *prr7prr9* mutants. The whole-plant context of the Framework Model then allowed us to test whether that metabolic change predicted an altered biomass in the mutants compared to wild-type plants. Mis-timed starch degradation largely (58-65% of the biomass reduction in experiments 1 and 2) or entirely (experiment 3) accounted for the mutants’ observed biomass, consistent with a previous, qualitative conclusion from other clock manipulations (Graf *et al.* 2010). Unused malate and fumarate accounted for the remaining biomass defect in experiments 1 and 2. These metabolite pools had previously been linked to clock gene function by machine-learning analysis (Grzegorczyk *et al.* 2015). The same mechanisms might affect the biomass growth of arrhythmic *prr5prr7prr9* mutants, which also accumulate these organic acids (Fukushima *et al.* 2009).

Where the *prr7prr9* double mutant plants mobilise the same absolute amount of starch per night as the wild-type plants, the most striking effect is an increased baseline starch level rather than a slower starch degradation rate, but the model explains this parsimoniously. The higher baseline starch level arises naturally if the plant’s starch metabolism in each unit of leaf mass is close to a steady state, where the absolute amount of starch degraded nightly equals the daily synthesis. Absolute starch synthesis per unit mass is also wild-type in *lsf1* control plants (Figure 3e), and this mutation is expected specifically to reduce the starch degradation rate. To degrade the same absolute amount of starch as wild-type plants but at a lower relative rate, the *lsf1* mutants must have a higher baseline starch level. The assumption of a lower relative degradation rate in *lsf1* is therefore functionally equivalent to the previous assumption of an altered ‘starch set point’ baseline level (Comparot-Moss *et al.* 2010, Scialdone *et al.* 2013). Why then do the mutant plants have lower total biomass, if they mobilise the same absolute amount of starch each night as the wild type in each unit of leaf mass (Figure 3e)? The explanation supported by the model is that the mutant plants accumulate large, unused starch pools as well as new biomass, whereas wild-type plants use the same photoassimilate supply to produce biomass more efficiently, leaving only a minimum of carbon in starch. As the plants continue to grow, the effect on biomass is cumulative: the leaf area that is not produced on one day results in a penalty on the plant’s total photosynthetic rate on every future day. Consistent with this mechanism, many parameter changes that reduced simulated starch levels at the end of the night also increased predicted biomass (Supporting Information Figure 7).

The biochemical mechanisms that connect clock output to starch degradation rate remain under investigation (Smith and Zeeman 2020). Neither of the candidate biochemical mechanisms for mis-regulation of malate and fumarate levels that we tested (PEPC activity and TDP level, Supplementary Figure 10) was obviously affected in the double mutants. Firstly, subtle changes might have significant, cumulative effects on final plant biomass and yet be below the resolution of our single-timepoint, biochemical assays. Secondly, a mathematical model cannot predict such biochemical mechanisms in much greater detail than has been tested in the available data. Including a richer model of central metabolism (Cheung *et al.* 2015) in the Framework Model could bring more of our metabolic data (Supporting Information Figure 3) to bear on this question. Until such causal mechanisms are determined, alternative explanations remain possible.

We assume that the altered starch levels cause the biomass defect in the *lsf1* mutants, because the LSF1 protein is located in the chloroplast and normally functions in starch degradation (Comparot-Moss *et al.* 2010, Schreier *et al.* 2019). The FMv2 simulations altered the starch levels of *prr7prr9* mutants by a very similar mechanism: removing PRR7 and PRR9 proteins from the clock sub-model predicted the clock-regulated slowing of the starch degradation rate and smaller plants as a result (as in Figure 3). Other models of starch dynamics have also represented a clock-regulated mechanism of this type (Pokhilko *et al.* 2014, Scialdone *et al.* 2013). However, an accurate model prediction is not a guarantee that the model is based on the correct explanation. We cannot infer the same causation as in *lsf1*, while the biochemical connection from the clock to starch regulation is unknown and cannot be directly tested in *prr7prr9*. We therefore conclude that the observed metabolic phenotypes <u>quantitatively account for</u> the clock mutants’ biomass growth defect, and that clock-regulated starch degradation is a feasible mechanism. In this scenario, the mechanisms of malate and fumarate mis-regulation in the clock mutants (and their altered carbon partitioning) are unclear, as noted above.

Alternatively or in addition, the clock might directly regulate growth rate, leading to mis-regulated growth in the mis-timed clock mutant plants. Slower growth in the *prr7prr9* double mutants might then leave unused photoassimilate to build up in various metabolite pools, including starch, malate and fumarate. Several lines of evidence suggest that such growth regulation is possible, though its mechanism is also unknown (Flis *et al.* 2019, Massonnet *et al.* 2011, Pullen *et al.* 2019). The gibberellin pathway was a candidate mechanism for this circadian effect, however our limited results were not obviously consistent with gibberellins as the mechanism for growth mis-regulation in *prr7prr9* (Supporting Information figure 11). The Framework Model could be adapted in future to distinguish these alternative directions of causation, as well as to test the mechanisms for a wide range of detailed behaviours that link the dynamics of light, circadian, metabolic and growth regulation.

Mugford et al. (2014) and Mengin et al. (2017), for example, suggest that the high sucrose observed after dawn in short photoperiods might accumulate due to the plant’s observed delay in resuming growth after a long night (and similarly after other perturbations that limit night-time sugar supply (Gibon *et al.* 2004, Moraes *et al.* 2019, Usadel *et al.* 2008)). This sucrose peak was proposed to increase AGPase activity allosterically (Supporting Information Figure 2a), and the FMv2 uses this outcome as the mechanism to increase carbon partitioning to starch under short photoperiods. Mugford et al. also showed that *fkf1* and perhaps *gi* mutants lacked this photoperiodic adjustment of partitioning (Mugford *et al.* 2014). The FMv2 already represents the key variables required to support future studies of this mechanism.

Whole-plant models based in molecular pathways are rare, because these studies remain challenging. Firstly, they require data of many types, ideally all acquired from the same plants, with calibration data at multiple scales that allow the models to use absolute units. Secondly, any environmental regulatory mechanism, such as the circadian clock, is sensitive to the experimental conditions, which can therefore contribute to variability in the data. Thirdly, there is often a balance to be struck between the specificity of an experimental manipulation (in this case the delayed timing of *prr7prr9)* and the magnitude of its effects, where an effect close in scale to stochastic, inter-plant variation can be laborious to reproduce but a more dramatic manipulation risks causing an unknown number of indirect effects, for example through carbon starvation. This work used broad experimental and modelling expertise, in organisations with dedicated facilities for plant systems biology, linked internationally by successive project awards (Supporting Information Figure 12). Accounting for the mild growth defect of *prr7prr9 via* particular biochemical mechanisms was at the limit of experimental tractability in our hands, due to the challenges noted above.

The approach has not been widely applied by Arabidopsis researchers, so we have considered how to facilitate its adoption. Users of the earlier FMv1 have been expert modellers, for example, so there has so far been little benefit from our investment in providing full access to FMv1 in user-friendly software with a graphical interface (Chew *et al.* 2014). A simpler, online simulator is therefore provided for non-experts to run the FMv2 (Supporting Information Figure 13). The scope of potential, future developments is broad. High-resolution phenotyping systems might soon resolve both physiological variation among individual plants and individual-specific, micro-environmental parameters, both in controlled environments and field conditions. The FMv2 could underpin a ‘digital twin’ approach to simulate each plant, which is closer in outlook to personalised medicine. Extensions of the Framework Model (Supporting Information Table 2) might test other critical rhythmic functions, such as photosynthesis, and address the nutrient and water limitations that prevail in field conditions. Even in our well-watered conditions, the *prr7prr9* clock mutants had consistently lower water content than wild-type plants (Supporting Information Figure 9k), suggesting another tractable effect that might be mediated by circadian misregulation of aquaporins and/or abscisic acid (Adams *et al.* 2018, Prado *et al.* 2019). Whole-organism physiology could also be understood (explained and predicted) quantitatively in other multicellular species (Le Novere 2015), for example using clock and metabolic models in animals and humans to understand body composition (Peek *et al.* 2013).

## Data

Access to new data is described in the Data and Model Availability section (Methods).

## Acknowledgements

Supported by European Commission FP7 collaborative project TiMet (contract 245143) to several authors, BBSRC FLIP fellowship BB/M017605 to A.H. and a BBSRC Institute Strategic Programme Grant BB/J004561/1 to the John Innes Centre. Financial support is gratefully acknowledged from the Swiss National Science Foundation (Grant 31003A_162555) and the University of Geneva to Teresa B. Fitzpatrick, from the Zürich–Basel Plant Science Centre Plant Fellows Programme (Marie Skłodowska-Curie Action Grant GA-2010-267243) and ETH Zurich to G.M.G and S.C.Z. For the purpose of open access, the authors have applied a CC-BY public copyright licence to any Author Accepted Manuscript version arising from this submission.

## Author Contributions

YHC, AS, MS and AJM designed the study. YHC, VM, AF, SM, AS and MS performed the main experiments and analysed the experimental data. YHC and DDS performed the modelling and analysed the simulation results. DDS, MM, TBF, GMG and SZ designed, performed and analysed follow-up studies. AH tested models and developed the online simulator. YHC, DDS and AJM wrote the paper with input from all authors.

## Methods

### Experimental methods

#### Plant materials and growth conditions

*Arabidopsis thaliana* of the Columbia (Col-0) accession, *prr7-3/prr9-1* (Nakamichi *et al.* 2007) and *lsf1-1* (Comparot-Moss *et al.* 2010) were used in this study. Seeds were first sown on half strength Murashige and Skoog (MS) solution and stratified in darkness at 4°C for 5 days before being exposed to white light at the desired photoperiod and temperature. Four-day-old seedlings were then transferred to soil containing Levington seed and modular compost (plus sand). The growth and treatment conditions for each experiment are shown in the figure legends. For the experiment in Figure 1d and Supporting Information Figure 3 only, seeds were sown on wet soil in pots and transferred directly to experimental conditions. Plants were thinned after a week and treated with nematodes after two weeks as a biological pest control.

#### Leaf number and plant assay

The total number of leaves (including the cotyledons) was recorded every 3-4 days from seedling emergence. Only leaves exceeding 1 mm^2^ in size (by eye) were considered in the total leaf count. Plants were harvested for biomass at different time points and for metabolite measurement at 3 weeks (Supporting Information Figure 3) and 4 weeks (other data). For metabolite measurement, rosettes were harvested and immediately submerged in liquid nitrogen, half an hour before lights off (end of day, ED) or lights on (end of night, EN) and stored at −80°C until extraction. For dry biomass, dissected plants were oven-dried at 80°C for 7 days. Area analysis was conducted using ImageJ (Schneider *et al.* 2012). Each image was first processed with colour thresholding to isolate the green region, which was next converted into binary format. The area was then determined using the Analyze Particles tool.

#### Gas exchange measurement

An EGM-4 Environmental Gas Monitor for CO_2_ (PP Systems, US) was used for CO_2_ flux measurement. A Plexiglass cylindrical chamber (12 cm in diameter x 3 cm sealed height, with a 6 cm tall support) was used (Supporting Information Figure 5f). Rubber rings around the lid and the hole for the pot ensured an airtight seal. The chamber was connected to the EGM-4 with two butyl tubes for closed-loop measurement.

Each individual measurement was taken by placing an individual plant pot in the chamber for approximately 60 seconds, during which the EGM-4 recorded CO_2_ concentration (μmol mol^-1^ or ppm) every 4.6 seconds. We covered the soil surface of the pots with black opaque plastic, leaving only a small hole in the middle for the plants. Plants were measured when they were 37 days old. Dark respiration was measured one hour before lights-on while daytime assimilation was measured one hour before lights-off.

CO_2_ enrichment of the atmosphere in the growth chambers due to the experimenters’ breathing was avoided by using a breath-scrubbing device during measurement. Hourly CO_2_ concentration at leaf level was also monitored by connecting the EGM-4 to a computer for automated data logging. The average hourly CO_2_ level was used as input to the model.

#### Extraction and determination of metabolite content

Rosettes were harvested as described above and ground in liquid nitrogen. Key results for 28-day-old plants are presented here; further results from the same experiments are available in the shared data files (see below). Around 20mg of ground material was aliquoted in screw-cap tubes (Micronic). Ethanolic extraction was performed using 80% ethanol v/v with 10mM MES (pH 5.9) and 50% ethanol v/v with 10mM MES (pH 5.9). During extraction, the successive supernatants obtained were combined into 96-deep well plates. The supernatant was used for spectrophotometric determination of chlorophylls, soluble carbohydrates, amino acids and organic acids as described (Arrivault *et al.* 2009). The pellet remaining after the ethanolic extraction was used for the determination of starch and total protein content as described (Pyl *et al.* 2012).

For Supporting Information Figure 10 (a-c), PEPC activity and incorporation of labelled ^14^CO_2_ into metabolite pools was measured as previously described for whole-plant labelling (Kölling *et al.* 2013, Kölling *et al.* 2015) with minor modifications. Near the end of the night (ZT20-21), 28-day-old plants were transferred to a custom built, sealed chamber in the dark. Each replicate consisted of the aerial parts of a single plant. Labelling was initiated through the release of ^14^CO_2_ (150 μCi) from NaH^14^CO_3_ with the addition of lactic acid. After 1 h, the plants were harvested into 5 ml 80% (v/v) ethanol and incubated for 15 min at 80°C. Following homogenisation, the soluble fraction was collected by centrifugation (2,400*g*, 12 min), pooled with four sequential 1-ml washes (50% ethanol, 20% ethanol, water, and then 80% ethanol) of the insoluble fraction pellet. The soluble fraction was concentrated under vacuum. A water-soluble sub-fraction was collected by dissolving the near-dry exsiccate in 2 ml water, while the remainder was dissolved in 2 ml 98% ethanol. Basic, acidic, and neutral fractions were separated from the water-soluble fraction by ion-exchange chromatography as described (Quick *et al.* 1989). Partitioning into protein was measured as described (Kölling *et al.* 2013). Isotope incorporation into each fraction was measured by liquid scintillation counting.

For Supporting Information Figure 10 (d-f), the extraction of B1 vitamers was performed using 100 mg of plant tissue homogenised in 200 μL of 1% (v/v) trichloroacetic acid. The mixture was centrifuged for 10 min at 10000 g and the supernatant decanted. B1 vitamers were derivatized and quantified by an HPLC method as described (Moulin *et al.* 2013). The data were normalized to the tissue fresh weight.

### Modelling methods

Development of the FMv2 in Matlab (Mathworks, Cambridge, UK), model equations, experimental data for model calibration and simulation procedures are described in the Supporting Information Methods section.

### Data and model availability

The data are shared collectively as a static Snapshot, formatted as a Research Object and structured according to the standard ISA hierarchy, on FAIRDOMHub.org (doi: 10.15490/FAIRDOMHUB.1.INVESTIGATION.123.??)[to be added in revision], on the Zenodo repository (doi: 10.5281/zenodo.??)[to be added in revision] and on the University of Edinburgh DataShare (doi: to be added in revision??). Within the Snapshot, the model is linked from the GitHub repository. The live, updatable resources are also public and structured on FAIRDOMHub.org, as https://fairdomhub.org/investigations/123 [note: this resource is accessible now]. The Open Data include results that are not analysed here, including metabolite levels at 21 days after sowing, and results from the *pgm* starch mutant in experiments 2 and 3 and the *lhy;cca1* clock double mutant in experiment 3 and the dark assimilation study. The data should be cited using the relevant DOI, for example as:

Chew, Y.H. et al. (2022). The Arabidopsis Framework Model version 2 predicts the organism-level effects of circadian clock gene mis-regulation [Data set]. Zenodo. https://doi.org/10.5281/zenodo.??[to be added in revision]

The luciferase reporter gene assays for circadian period of Col-0 and *prr7prr9* seedlings, with and without exogenous GA, are available from the BioDare repository as BioDare experiment ID 3838: choose https://biodare.ed.ac.uk/experiment (“Browse Public Resources” on the Login screen), then https://biodare.ed.ac.uk/robust/ShowExperiment.action?experimentId=3838, “Effects of GA on clock in WT and prr9prr7”. Raw and processed data, metadata, period analysis and summary statistics are available.

A simple, online simulator allows non-experts to run the FMv2 for wild type and *prr7prr9* in multiple conditions, with a public, web browser interface at http://turnip.bio.ed.ac.uk/fm/. (Supporting Information Figure 13).

## Supporting Information Tables

**Supporting Information Table 1:**
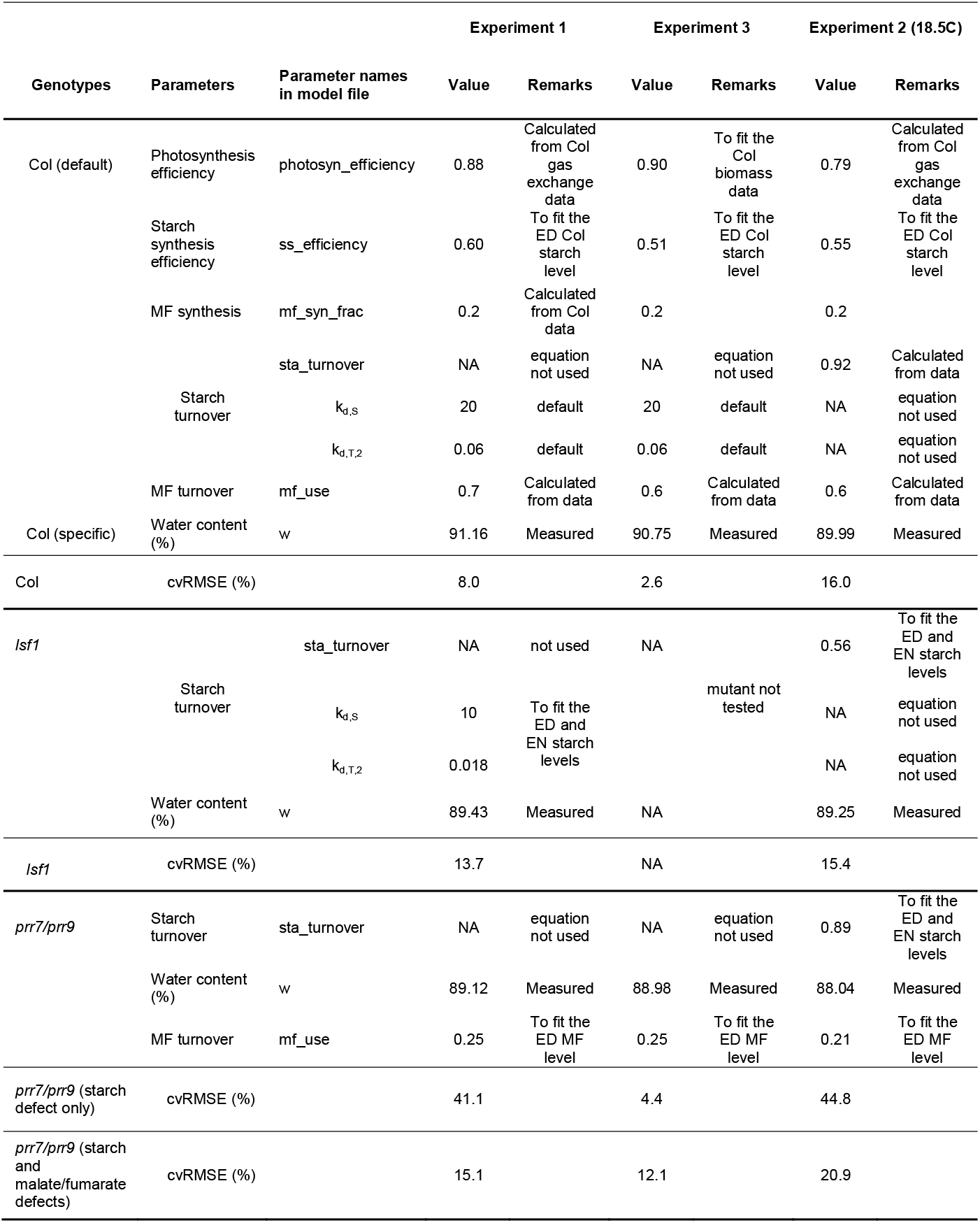
Parameter and cvRMSE values for all studies and genotypes.

**Supporting Information Table 2.**
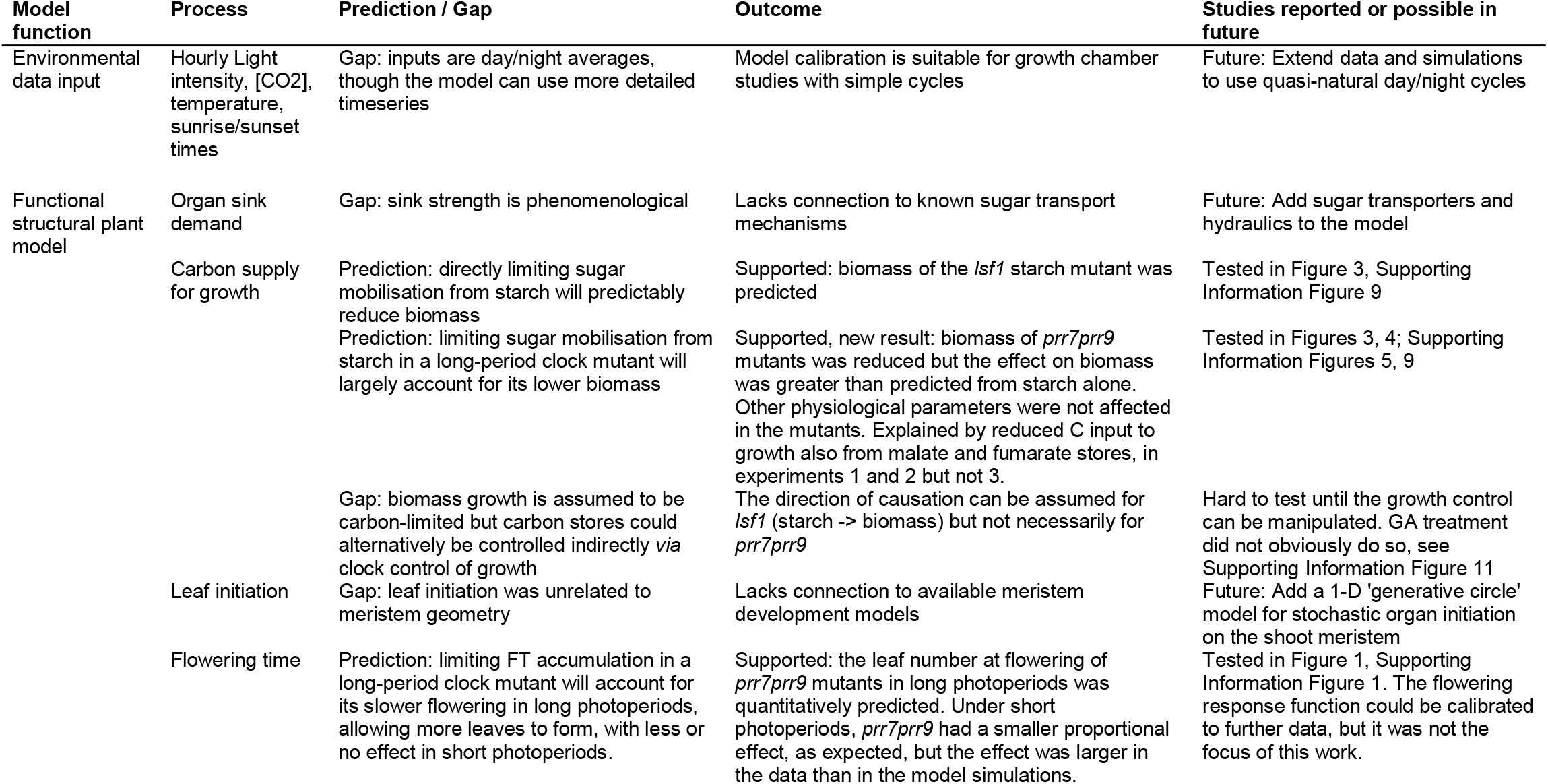
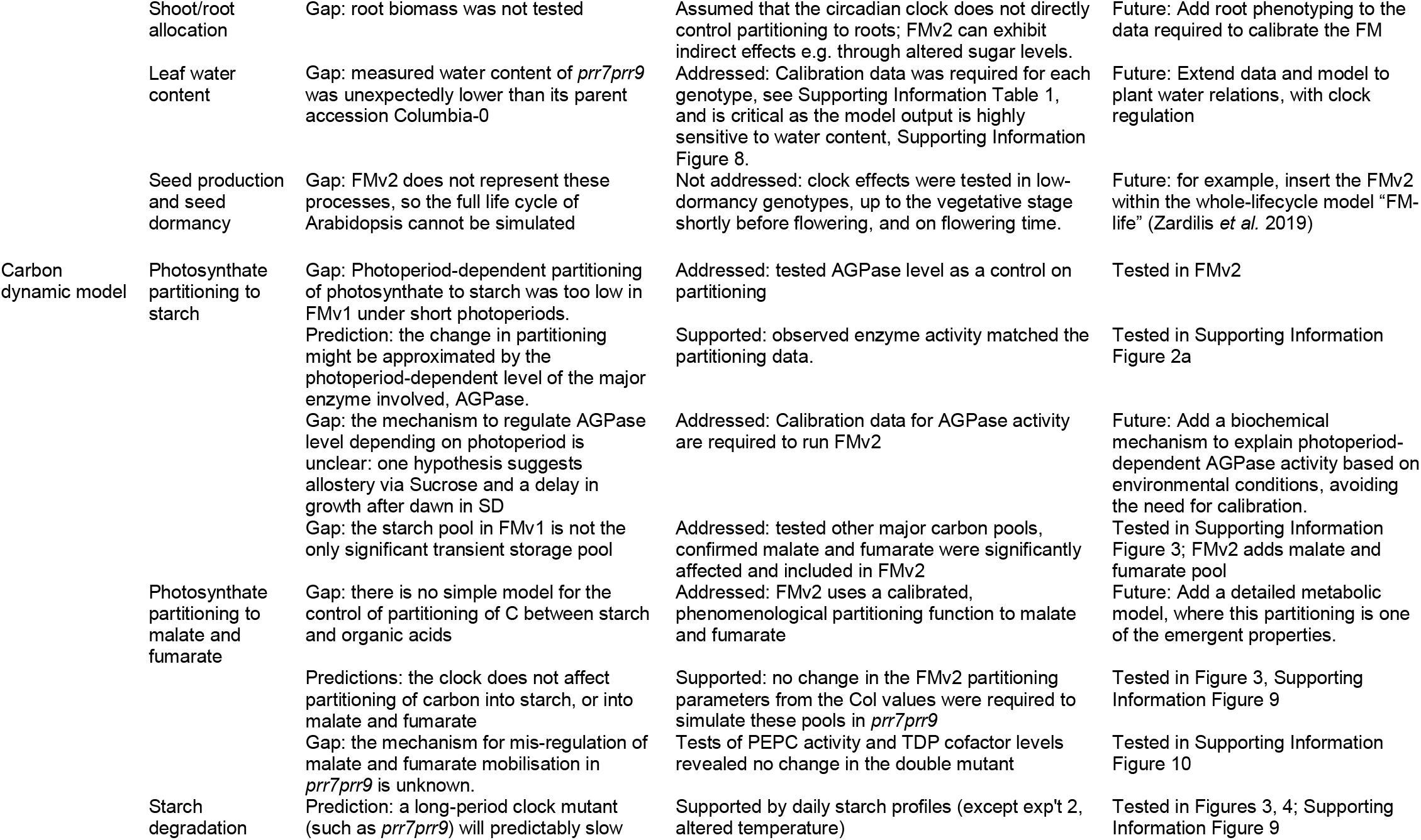
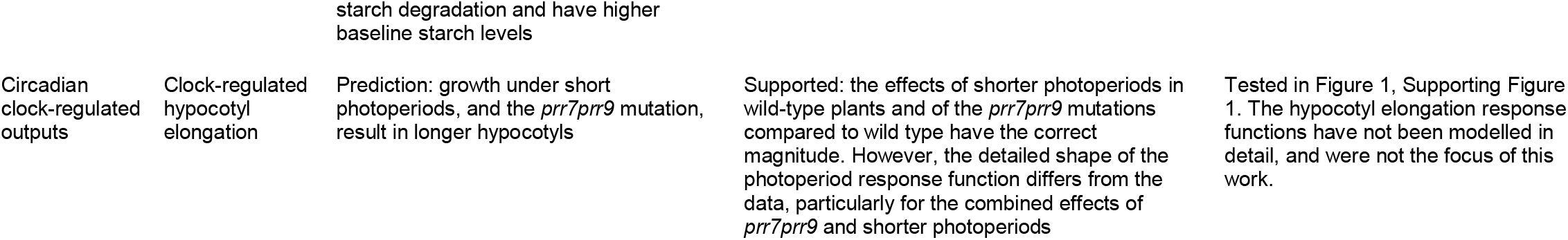
Outcomes from the FMv2: predictions, gaps and future studies. The results described in this paper are summarised, with potential future studies, and some aspects of the FMv2 that were not tested here.

### Supporting Information Figure Legends

**Supporting Information Figure 1:**
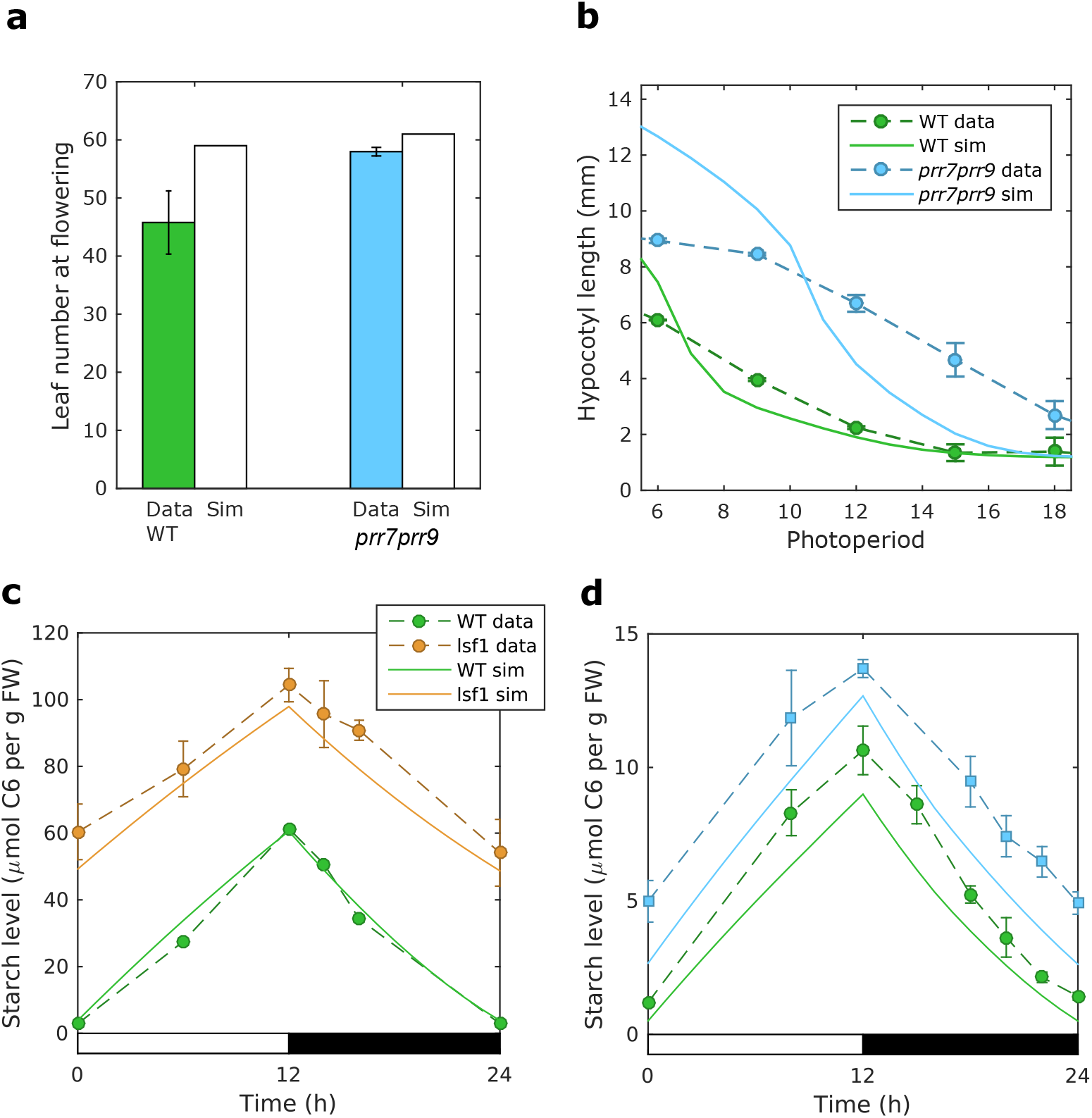
Simulation of clock outputs. (a) Measured leaf number under short photoperiods for Col WT (green) and *prr7prr9* (blue) (Nakamichi *et al.* 2007), compared to simulation (white); (b) Measured hypocotyl elongation in multiple photoperiods (Niwa *et al.* 2009), compared to simulation (solid lines); (c) starch levels in Col wild type plants (WT, green) and *lsf1* mutants (orange) under 12L:12D cycles (Comparot-Moss *et al.* 2010), compared to model simulations (solid lines); (d) Starch levels in Col wild type plants (WT, green) and *prr7prr9* mutants (blue) under 12L:12D, compared to model simulations (lines); as in Fig 3a, plotted here as absolute values. Data are mean ± SEM (n = 6). Temperature= 20 °C; light = 190 μmol/m^2^/s; photoperiod = 12h light: 12h dark.

**Supporting Information Figure 2:**
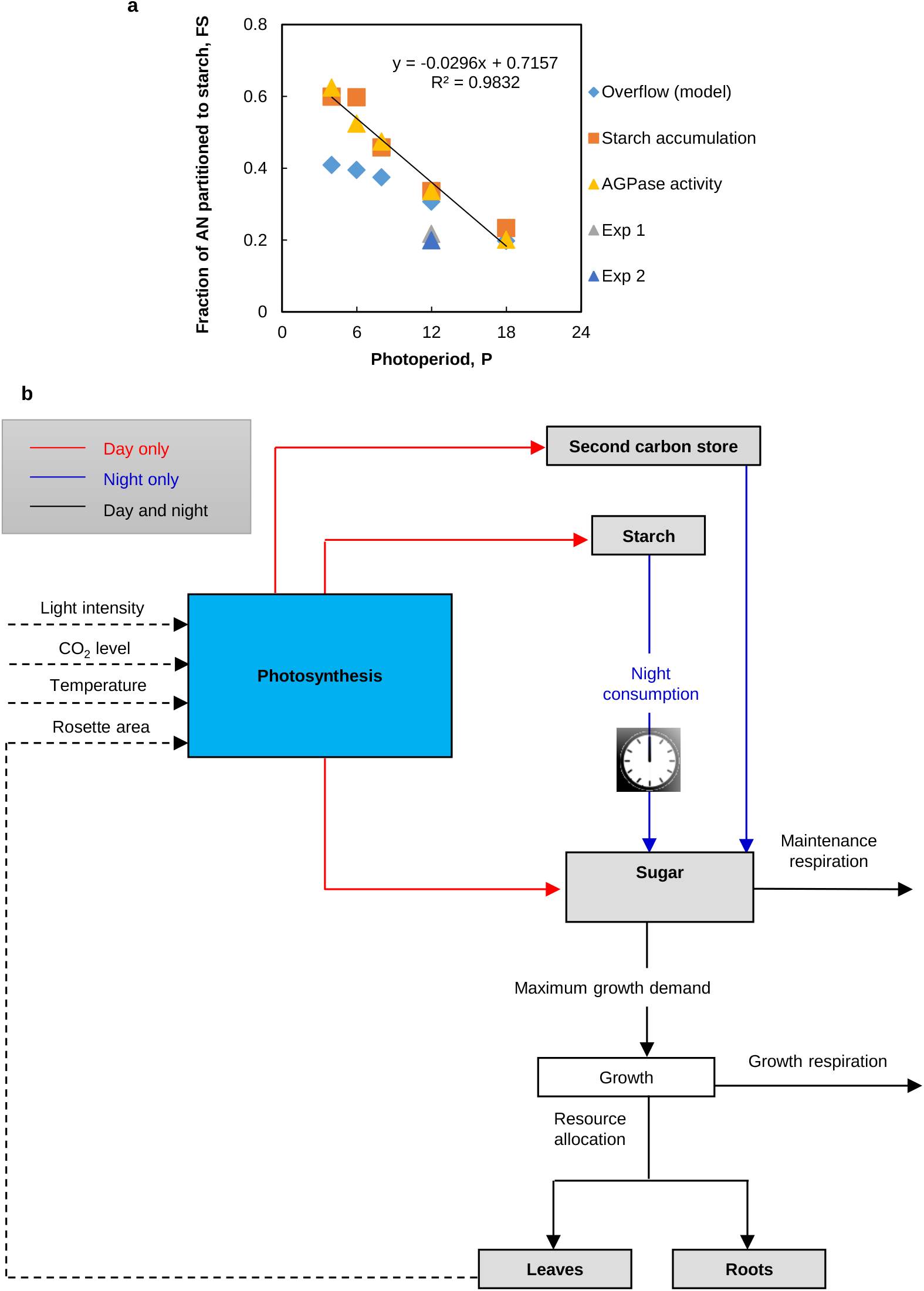
Updating the Carbon Dynamic Model (CDM). (a) The fraction of net assimilate partitioned to starch in plants grown under different photoperiods, as simulated in FMv1 using the ‘overflow’ concept (Overflow, blue diamonds), calculated based on measured starch levels (Gibon *et al.* 2009) (starch accumulation, orange) and calculated based on measured AGPase activity normalised to the value in 12:12D (AGPase activity, yellow). The regression line with equation inset is for the AGPase activity series. Triangles show the values calculated from measured starch levels in experiments under 12L:12D (Experiment 1, grey; Experiment 2, blue; Experiment 3 not shown, 7% lower than Experiment 2), reflected in the value of model parameter ss_efficiency (Supporting Information Table 1). (b) Schematic of the updated Carbon Dynamic Model (CDM). The second carbon store represents the total amount of malate and fumarate. The clock symbol represents the regulation of the rate of starch consumption by the circadian clock model. Red/blue arrows indicate day-/night-specific processes. Dashed arrows indicate information or feedback input.

**Supporting Information Figure 3:**
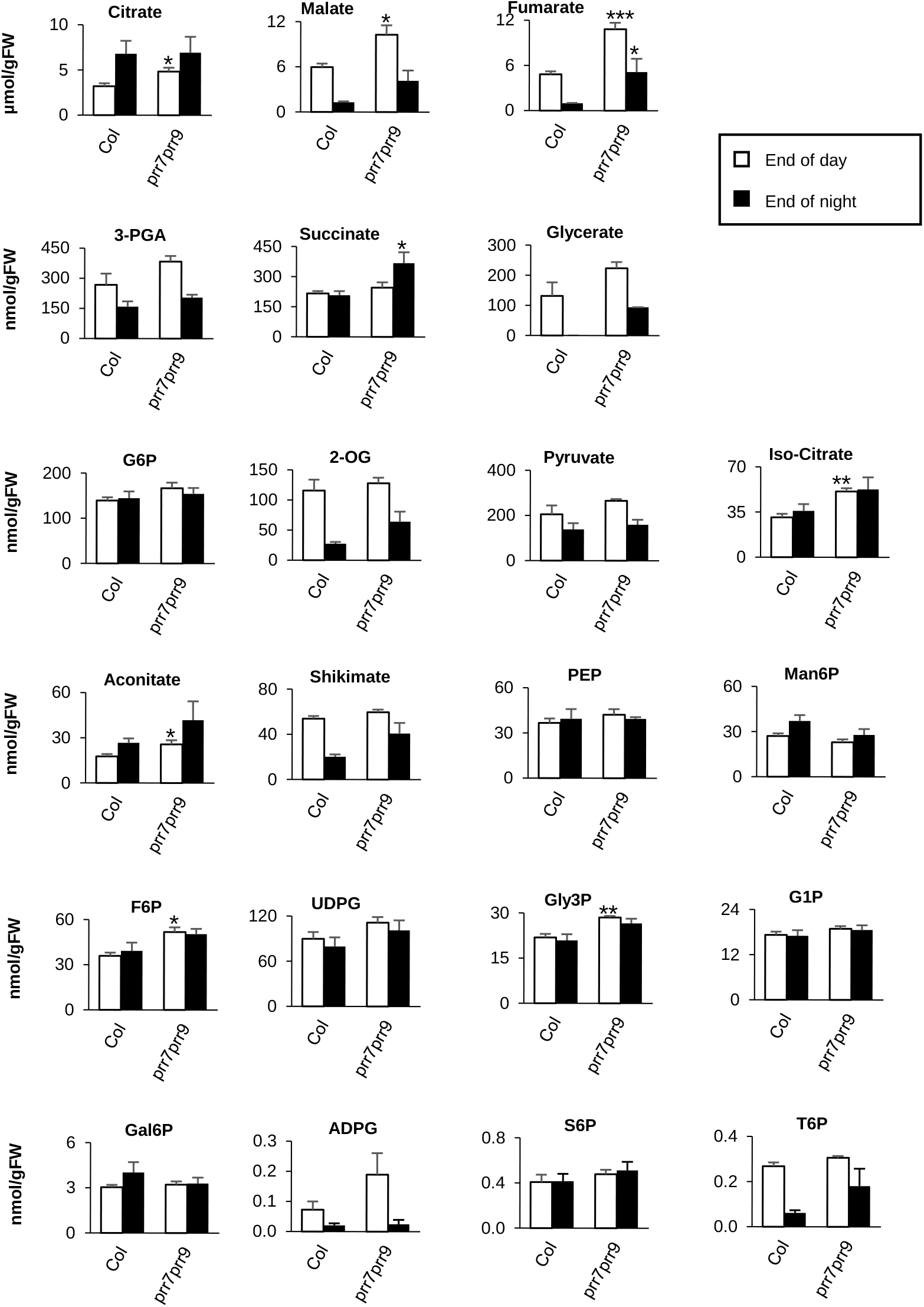
Primary metabolites for Col and *prr7prr9* mutant plants measured at the end of day and the end of night. The results of metabolite analysis are given as the mean ± SEM (n = 4). Each sample consisted of 5-7 pooled plants harvested at the end of day (open bars) or end of night (filled bars). Temperature = 20°C during the day and 18°C during the night; light = 160 μmol/m2/s; photoperiod = 12 h light: 12 h dark. The t-test compared between Col and prr7prr9 (* p < 0.05; ** p < 0.005; *** p < 0.001). Note y-axis scales, and units given for citrate, malate, and fumarate are μmol/gFW, while the units given for the remaining metabolites are nmol/gFW. The measurement of glycerate failed in the Col samples for the end of the night.

**Supporting Information Figure 4:**
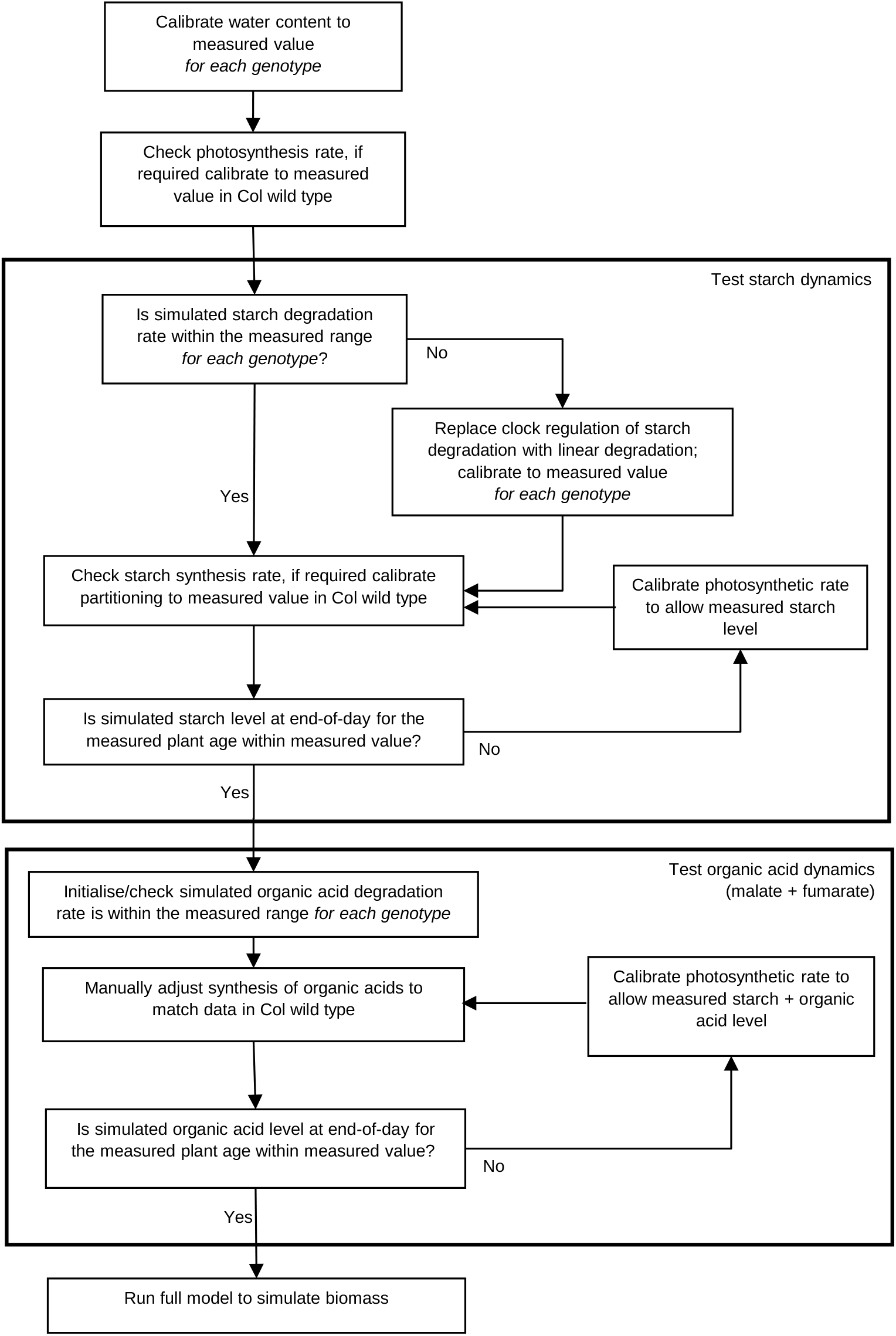
Flow diagram of parameter calibration. Parameters that could be directly or indirectly measured were adjusted in the illustrated sequence, to capture measured carbon dynamics and metabolite levels at specific time points. Once these were achieved, the model was simulated using the determined parameters to generate predictions for plant biomass.

**Supporting Information Figure 5:**
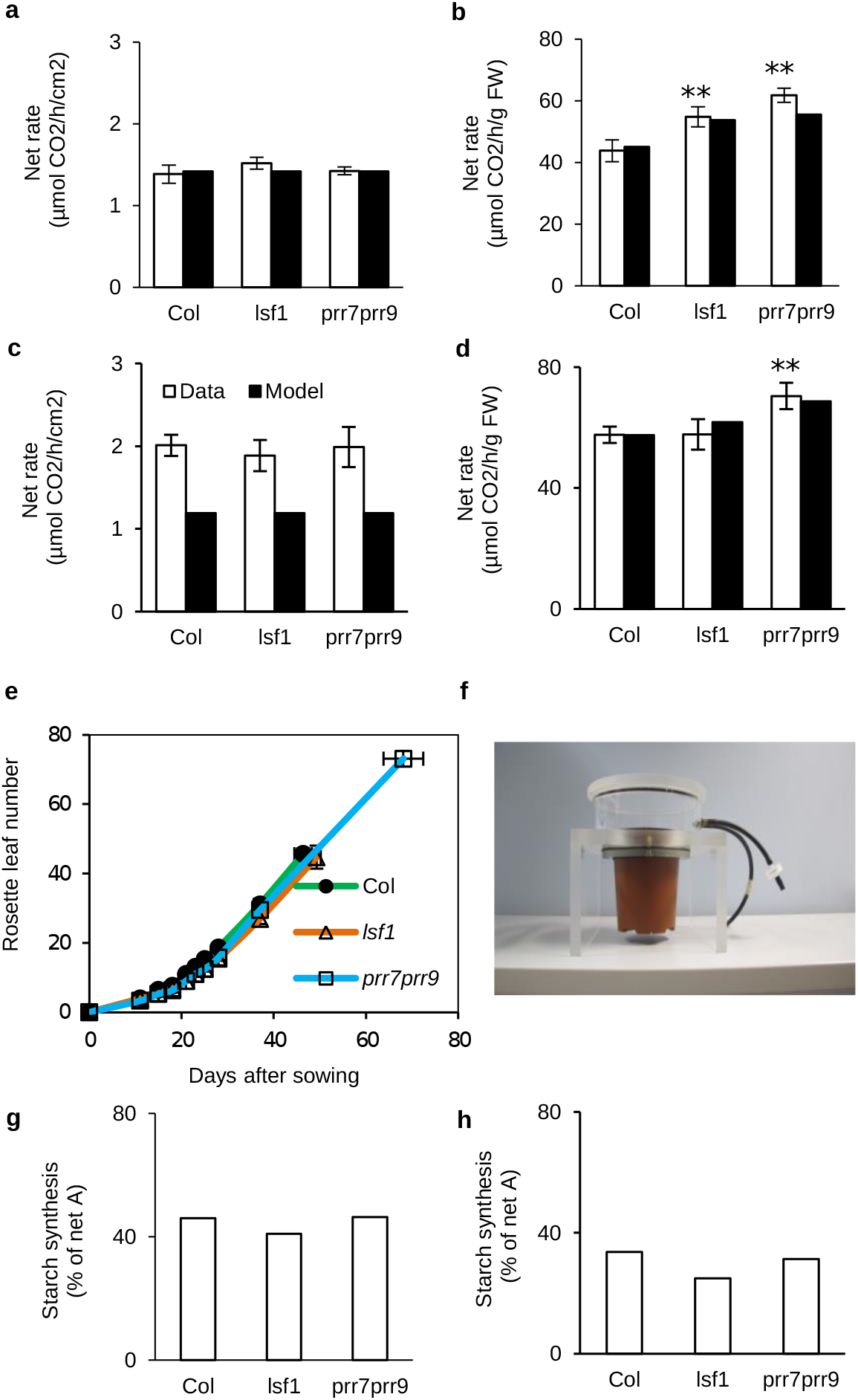
Gas exchange measurement and leaf number of different mutants. Net assimilation rate for Col, *lsf1* and *prr7prr9* in Experiment 1 (a, b) and Experiment 2 (c, d) expressed per unit rosette area (left column) and per gram fresh weight (right column). Data are shown as white bars while model simulations are shown as black bars. Data are given as mean ± SD (n = 5 plants). There were no significant genotypic differences for net assimilation per unit area, thus similar rates were used in model simulations for all genotypes. However, net assimilation per unit fresh weight was significantly higher in *lsf1* and *prr7prr9* (** p < 0.005). Taking genotypic differences in water content into account was sufficient for the model to reproduce these results. (e) Rosette leaf number until flowering for Experiment 1. There was no significant difference in the leaf number at flowering for *lsf1*, while *prr7prr9* was late-flowering as expected. (f) The Plexiglass chamber used for gas exchange measurement. Starch synthesis as a percentage of net assimilation rate for Experiment 1 (g) and Experiment 2 (h).

**Supporting Information Figure 6:**
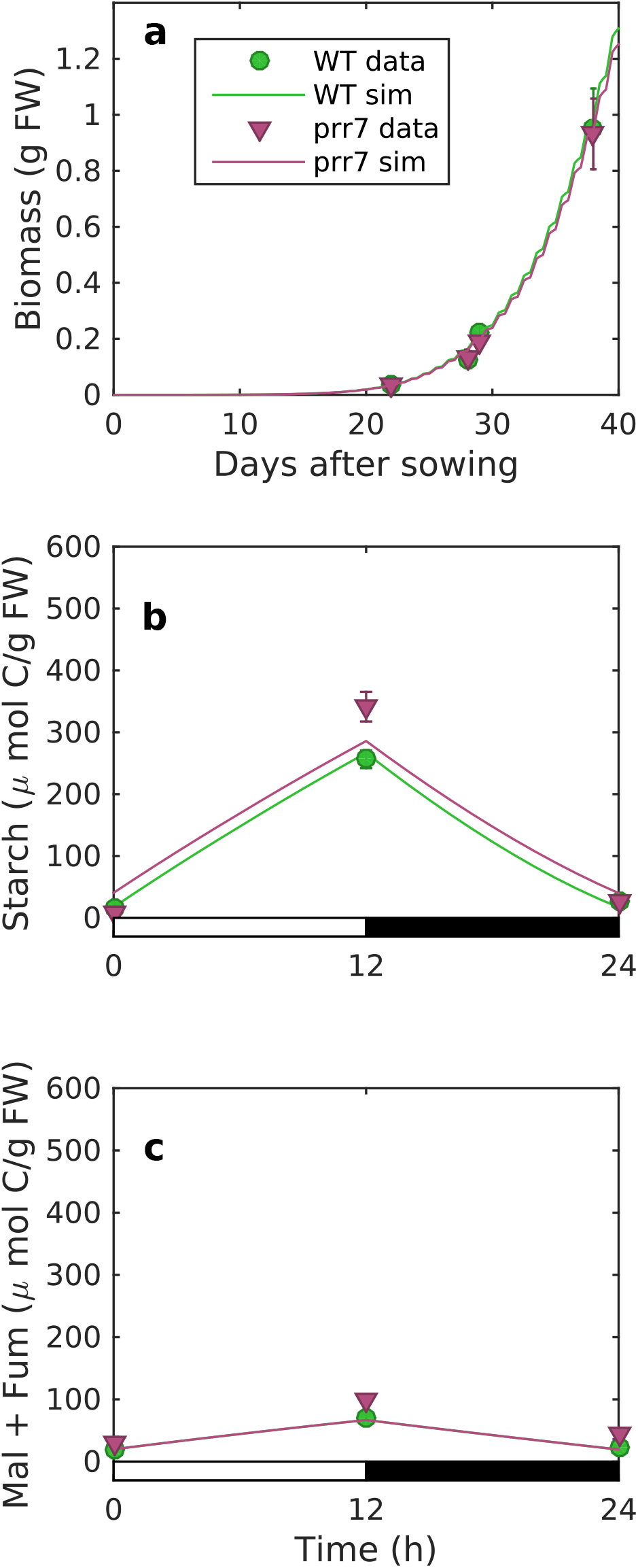
Biomass and carbon status of *prr7* mutants. Measured (symbols) and simulated (lines) fresh weight (a), starch level (b), and sum of malate and fumarate (c) for *prr7* single-mutant plants (purple) compared to wild type Col (green) in Experiment 1. Mean +/− SEM, n=5 for biomass; n=3 for metabolites, where each sample pooled 3 plants. Data for Col are identical to Figure 3c,3e,3g.

**Supporting Information Figure 7:**
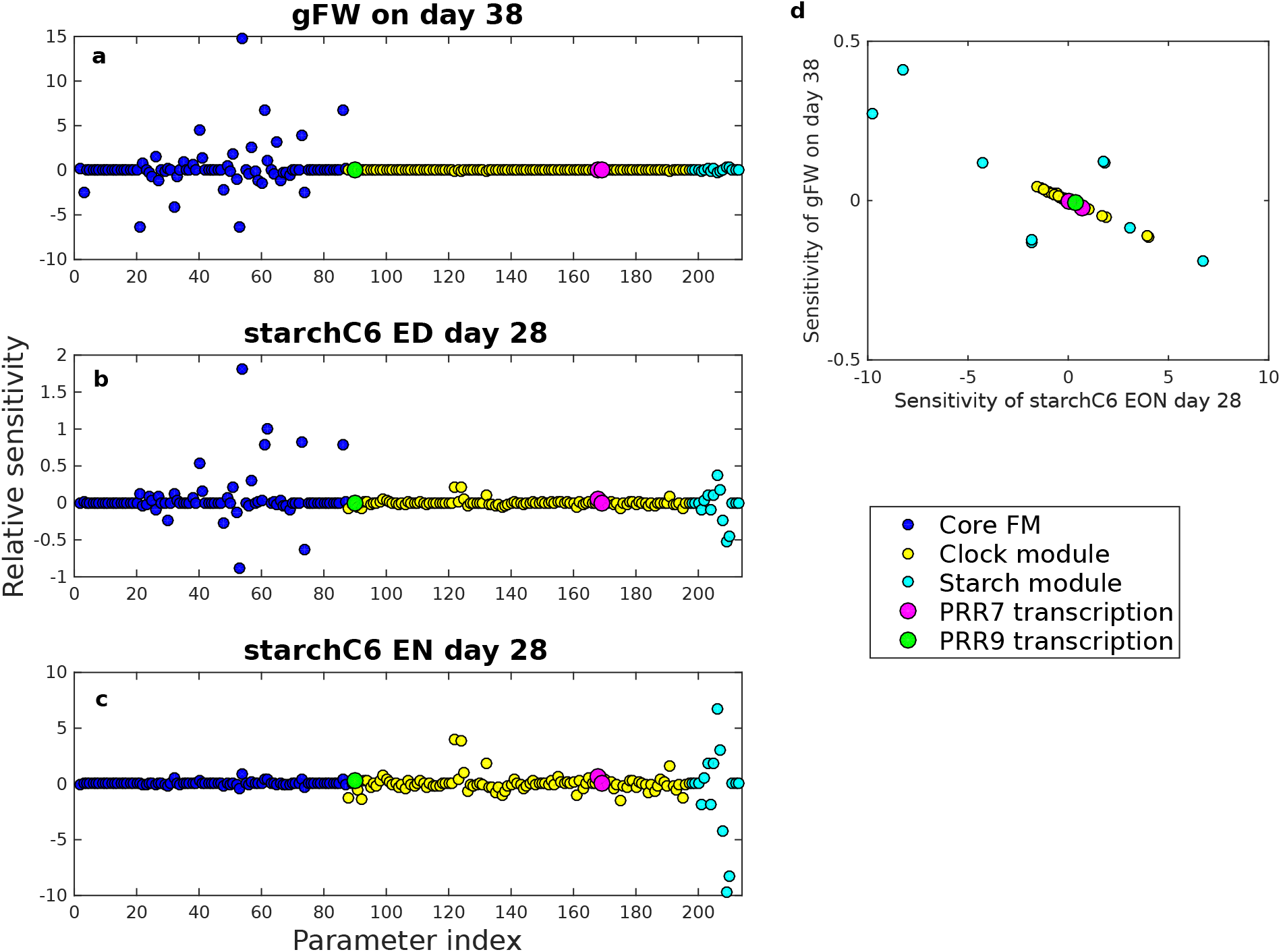
Parameter sensitivity overview. Relative sensitivity of model outputs for each model parameter, coloured by the cognate sub-model (see legend). Sensitivities were calculated by simulating the model under 1% perturbations of each parameter in turn. (a) Fresh Weight (FW); most sensitive parameter is s_elec_, associated with electron transport, (b) starch level at end of day (ED); most sensitive parameter is s_elec_ and (c) at end of night (EN); most sensitive parameter is k_dT1_, the degradation rate of a putative inhibitor of starch turnover. Clock parameters mutated to simulate *prr7prr9* double mutants are highlighted (see legend). (d) Comparison of sensitivity of FW and EON starch for parameters clock and starch module parameters, showing a predominant negative correlation: parameters that lower starch at the end of the night tend to increase fresh weight. Note that water content (w, a directly measured parameter) is not shown due to high sensitivity. Sensitivities to changes in water content were 11.3, −10.1, and −10.1 for gFW, ED starch, and EN starch, respectively (see also Supporting Information Figure 8).

**Supporting Information Figure 8:**
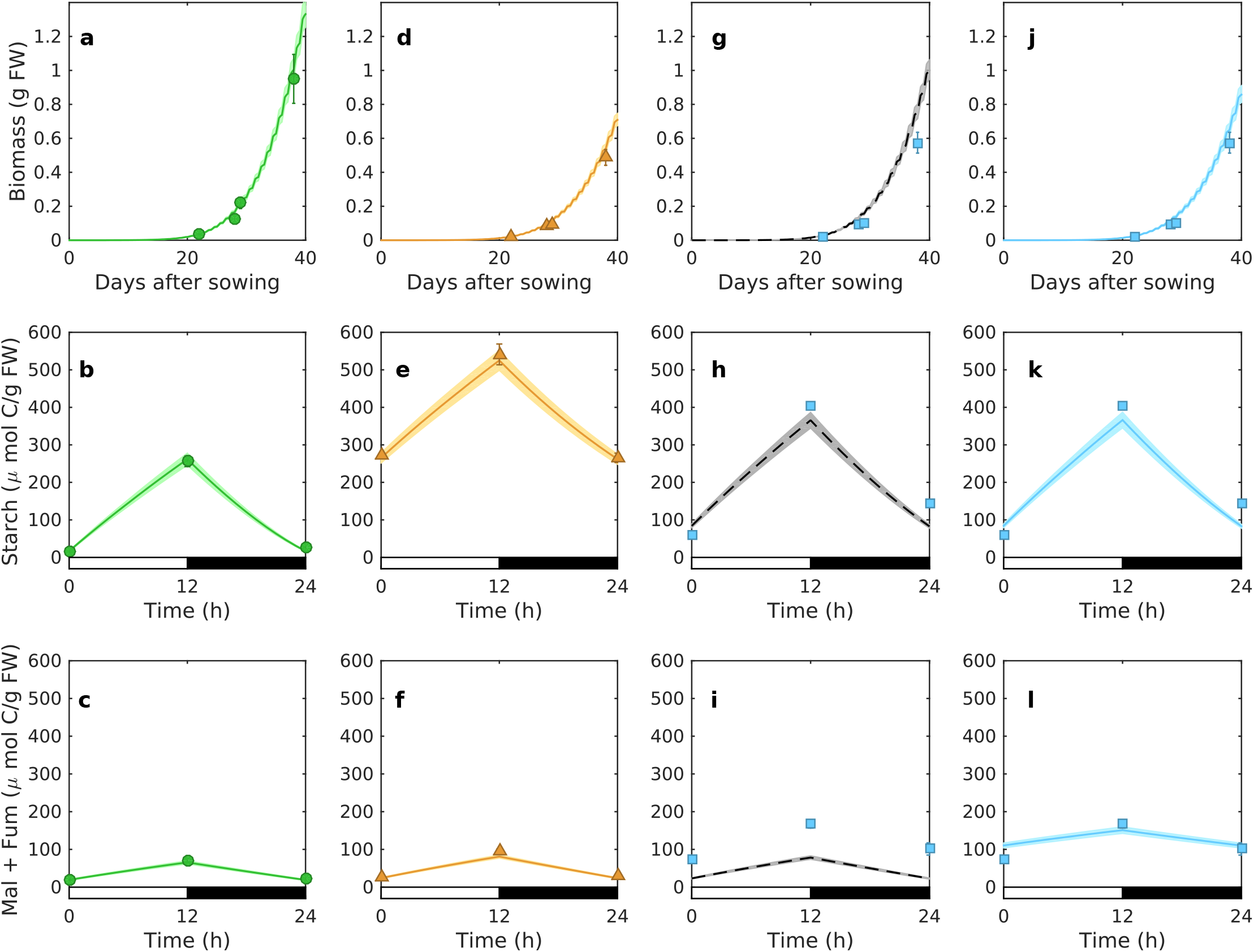
Model sensitivity to water content parameter. Simulation of biomass and major metabolites is shown using water content values plus and minus one standard deviation from the mean, for Experiment 1 (Figure 3). Model simulation (lines) and experimental data (symbols) of fresh weight (a,d,g,j), starch level (b,e,h,k) and the total level of malate and fumarate (c,f,i,l) for Col (a-c), *lsf1* (d-f) and *prr7prr9* (g-l). Dashed lines (g-i) are model simulation for *prr7prr9* that only considered starch defects, solid lines (j-l) are model simulations that included both starch defects and inefficient use of malate and fumarate. Shaded regions indicate the values spanned by simulating water contents plus and minus one standard deviation from the mean, for each genotype. Data are as in Figure 3: mean ± SD (n = 5 for biomass; n = 3 for metabolites with each sample consisting of 3 pooled plants). Temperature = 20.5°C; light=145 μmol/m2/s; photoperiod = 12 h light: 12 h dark; CO_2_ = 420 ppm.

**Supporting Information Figure 9:**
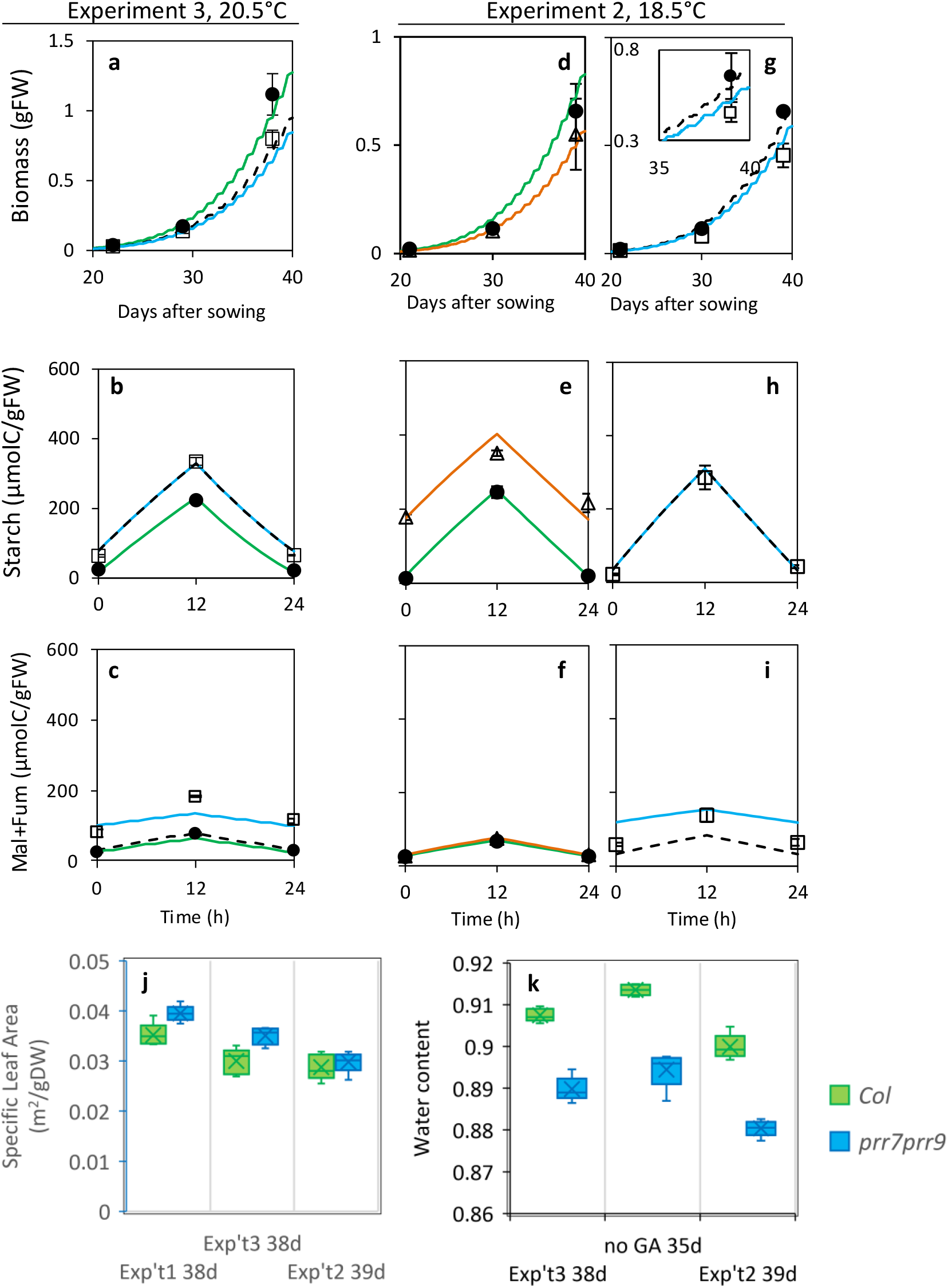
Repeated tests of biomass growth, starch and organic acid content. Experiment 1 (Figure 3) was repeated in experiment 3 (a-c) and at a lower temperature in experiment 2 (d-i). (ch) Data (symbols) and simulation (lines) of fresh weight (a,d,g), starch (b,e,h) and total malate and fumarate (c,f,i) for Col (circles, green), *lsf1* (triangles, orange) and *prr7prr9* plants (squares). Simulations of *prr7prr9* show a starch defect only (dashed black line) or both starch and malate+fumarate defects (solid blue). (g) Inset enlarges main panel around 39-day data point. Data show mean±SD; n=5 for biomass; n=3 for metabolites, where each sample pooled 3 plants. Y-axis scale for biomass plots differs between experiments 2 (d,g) and 3 (a). Starch and malate and fumarate data are plotted on the same scales as in Figure 3 (e-h). Growth conditions, 12L:12D with light intensity= 145 μmol/m^2^/s, temperature 20.5°C (a-c), 18.5°C (d-i); CO_2_=420 ppm. (j) The mean specific leaf area for *prr7prr9* plants is greater than for Col-0 (ANOVA p < 0.001). (k) *prr7prr9* plants have lower water content. j, k - Data are mean, n=5, error bar = SD, from the same plants as tested for biomass. Mean water content was measured using pooled plants in experiment 1, not shown.

**Supporting Information Figure 10:**
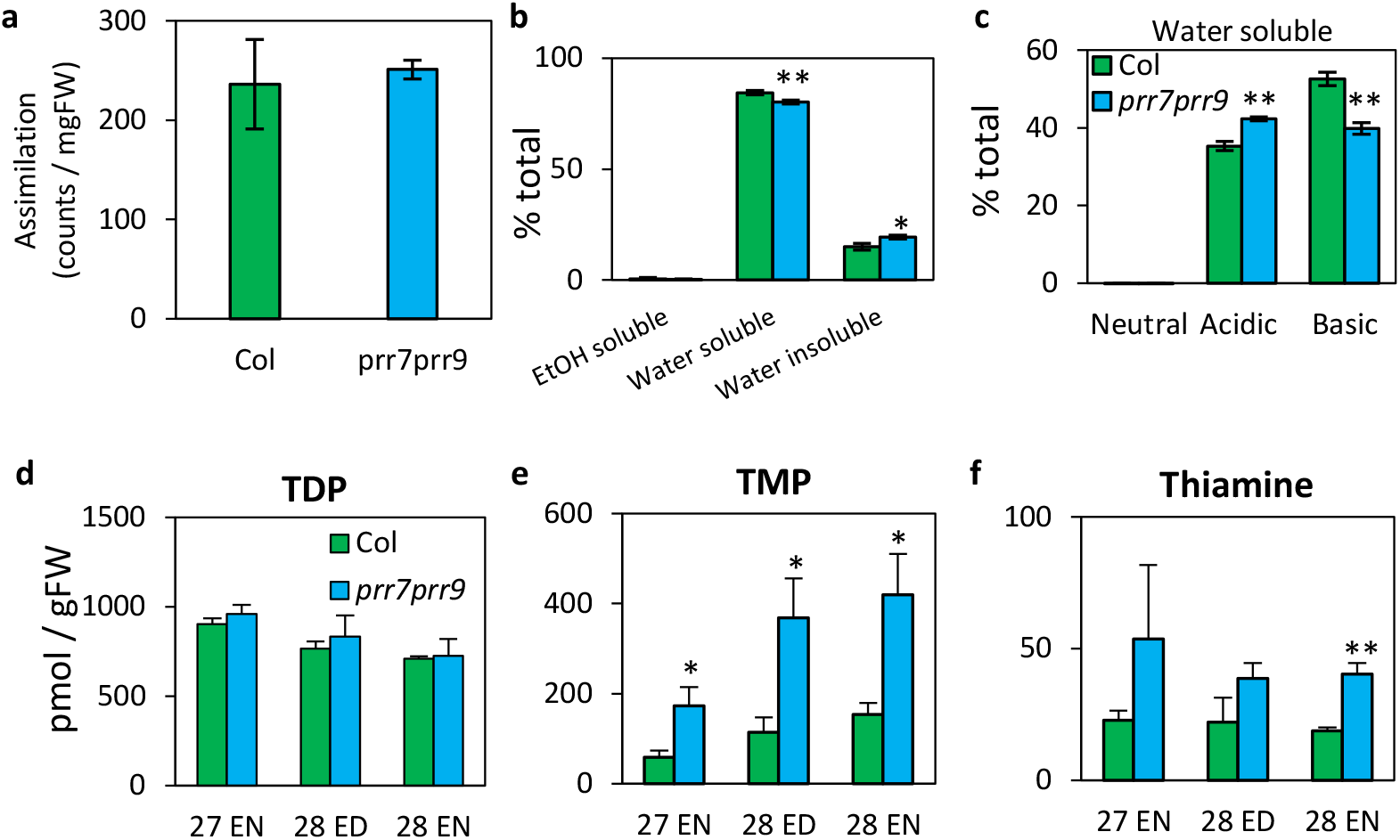
Testing direct mechanisms of malate and fumarate accumulation. Perturbations of organic acid synthesis in 28-day-old Col wild type and *prr7prr9* plants were tested directly by ^14^CO_2_ labelling in darkness before lights-on (a-c) and indirectly *via* the levels of thiamine-derived cofactors for metabolic enzymes (d-f). a) Carbon assimilation through PEP carboxylase was similar in Col and *prr7prr9;* b) percentage of assimilated carbon in ethanol soluble, water soluble and water insoluble fractions; c) partitioning of assimilated carbon within the water soluble fraction into neutral, acidic and basic fractions; d-f) HPLC analysis of thiamine-related compounds shows similar levels of TDP (d), but higher TMP (e) and thiamine (f) in *prr7prr9* samples compared to Col, in plant samples from experiment 3 (also analysed in Supporting Information Figures 9h, 9i). Data shown are mean ± SD, n = 3 (a-c) or n = 4 (d-f). The statistical significance of differences between Col and *prr7prr9* is indicated by t-test probability, * p<0.05, ** p<0.005. Temperature = 20-20.5°C; light=145-150 μmol/m^2^/s; photoperiod = 12 h light: 12 h dark; CO_2_ = 420 ppm.

**Supporting Information Figure 11:**
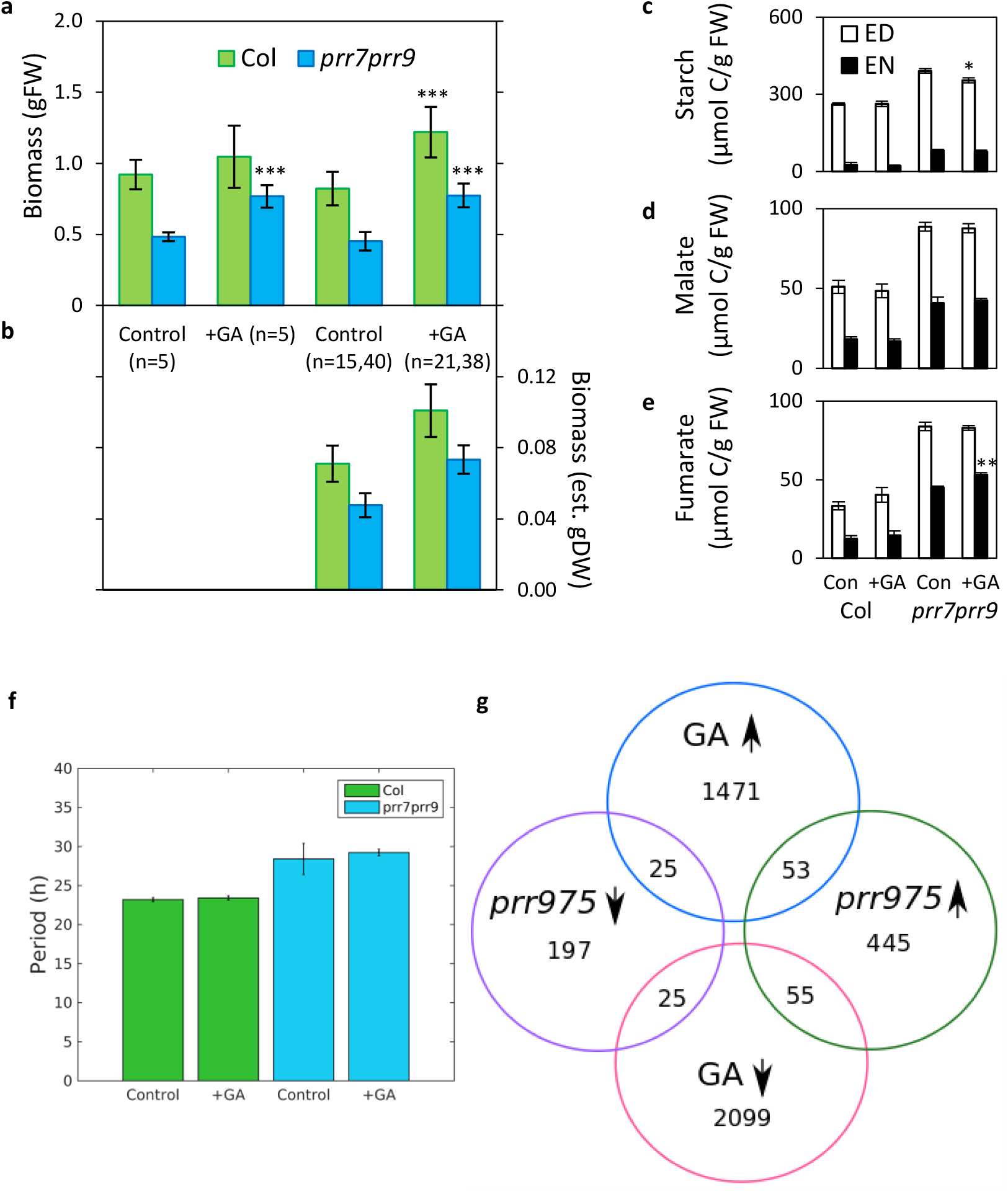
Testing gibberellin-stimulated growth for reversal of the clock-mediated carbon accumulation. a-e) Biomass and metabolite analysis of plants grown with or without GA treatment. a) fresh biomass at 35 days after sowing in the subset of plants tested for dry biomass (n=5) and in all plants (n as indicated). b) estimated dry biomass of all plants, calculated by applying the average water content measured in each n=5 subset to the fresh biomass data of the corresponding full set shown in a). c-e) metabolite analysis of Col-0 and *prr7prr9* plants at 28 days after sowing, for c) starch, d) malate and e) fumarate, at the end of the day (ED) or end of the night (EN). n=3 as in Figure 3 and Supporting Information Figure 9. All data are means, error bar = SD. t-test probability of a difference between GA-treated and control samples is marked *, p<0.05; **, p<0.01, ***, p<0.005. Comparisons of *prr7prr9* to Col are highly significant, p<0.001. f) GA treatment did not reduce the long circadian period of *prr7prr9* seedlings carrying the *CCR2:LUC* reporter gene, tested by *in vivo* imaging under constant light. g) The Venn diagram shows a minimal overlap between GA- and *PRR*-regulated transcripts. Lists of transcripts up- and down-regulated in the *prr5prr7prr9* triple mutant and after GA treatment were taken from (Nakamichi *et al.* 2009b) and (Bai *et al.* 2012), respectively. No consistent pattern was identified between the gene sets differentially expressed after GA treatment and in the *prr5prr7prr9* mutant.

**Supporting Information Figure 12:**
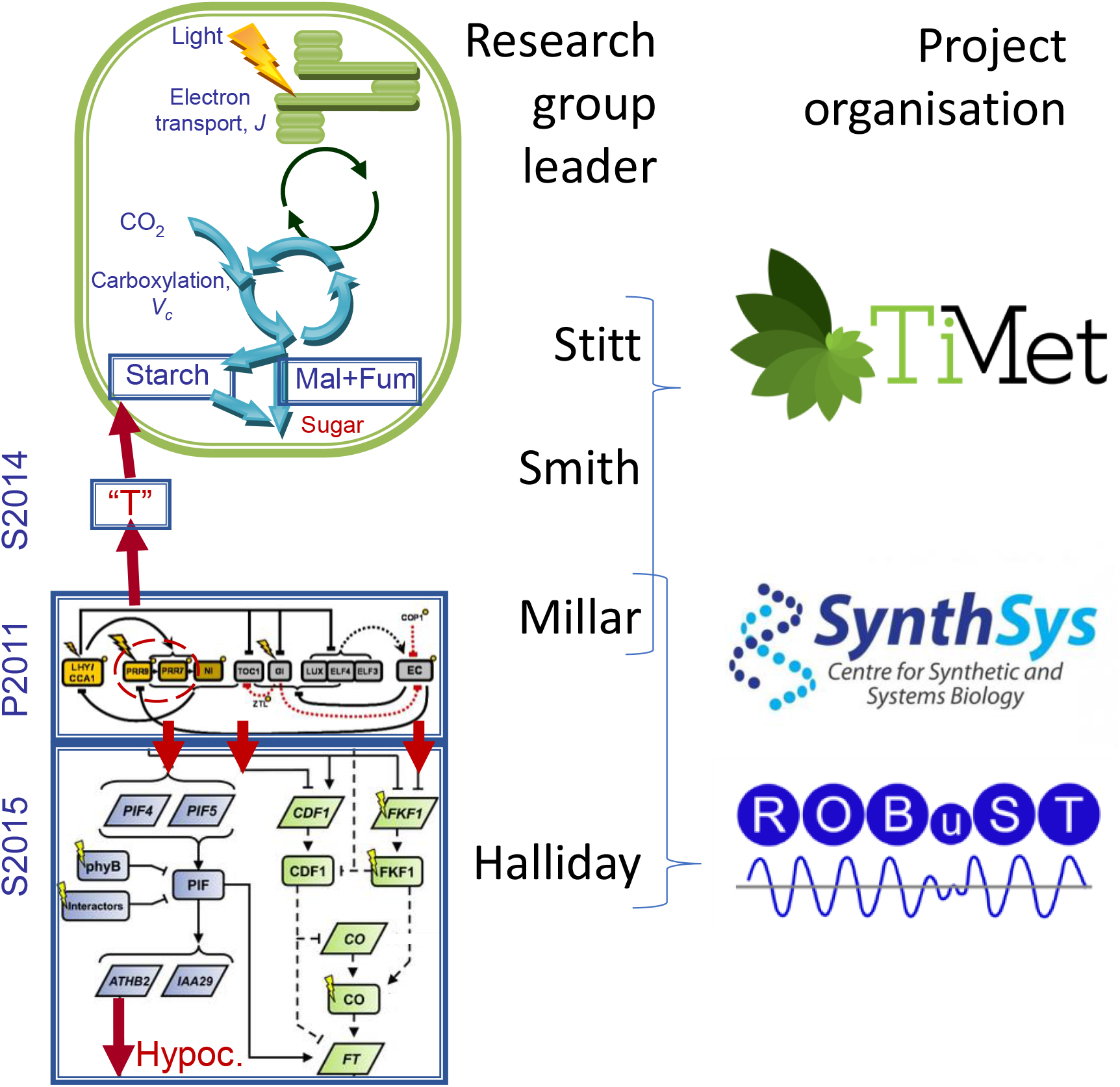
Provenance of the Arabidopsis Framework Model version 2 (FMv2). Model components that are updated in FMv2 are bounded in blue rectangles, with abbreviated model names (left), reproduced from Figure 2. To indicate the main contributors to model development, research group leaders are shown (centre) close to their area of contribution, in metabolism (Stitt, Smith), the circadian clock (Millar) and the light-response pathways and physiology (Halliday). The Fitzpatrick and Zeeman groups tested specific hypotheses arising from FMv2 and are not shown here. The funded research organisations that brought these researchers together are represented by their logos (right): the €6M EU FP7-funded TiMet project (245143; Stitt, Smith, Millar and others; S2014 and FMv2 models), the £11M BBSRC/EPSRC-funded SynthSys centre (BB/ D019621; Millar and others; P2011 model) and the £6M BBSRC-funded ROBuST project (BB/F005237; Halliday, Millar and others; S2015 model). All the partners and projects shown had earlier contributed to the FMv1 model.

**Supporting Information Figure 13.**
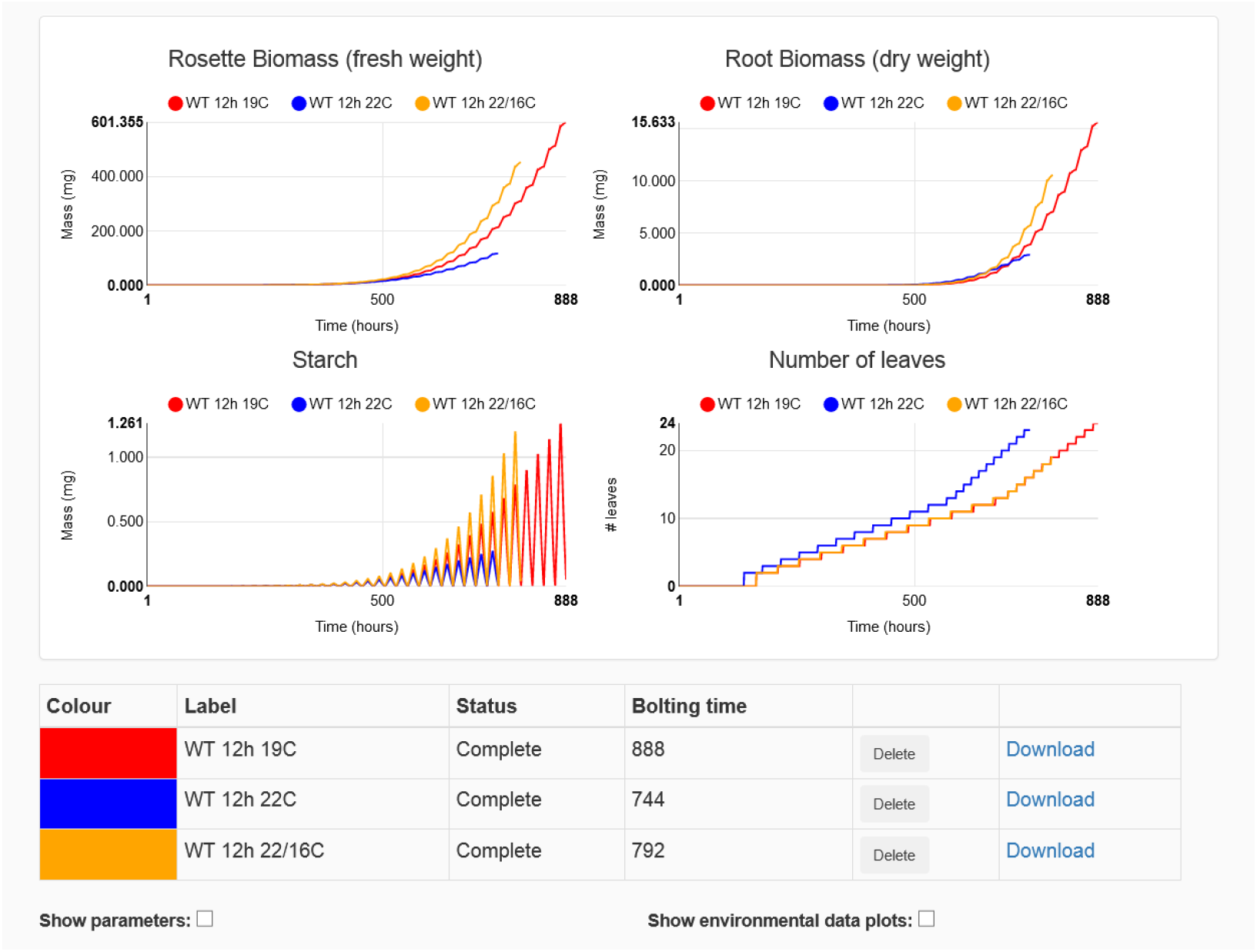
Online simulator for the Arabidopsis Framework Model version 2 (FMv2). Non-experts can simulate the FMv2 in a public, web browser interface at http://turnip.bio.ed.ac.uk/fm/. Simulations can select wild type or *prr7prr9* genotype, control environmental light, temperature and CO_2_ levels, and superimpose the results of successive simulations. The example shown compares wild-type plants simulated under 12L:12D cycles at constant 19°C (red traces) or 22°C (blue) with 22°C daytime temperature and 16°C nights (amber). The plotted model outputs are fixed: rosette fresh biomass and root dry biomass, starch and leaf number. Each plot ends with flowering. The hours to flowering (bolting time) are displayed, and timeseries outputs are available to download.

