## Supporting Information text for "The Arabidopsis Framework Model version 2 predicts the organism-level effects of circadian clock gene mis-regulation"

### Supplementary Information

#### Contents


1. Updating the circadian clock, starch, and photoperiod response models ... 1
   1.1 Photoperiod response model ... 1
   1.2 Circadian control of starch turnover ... 2
2. Revision of Starch synthesis ... 3
3. Addition of carbon pool for malate and fumarate ... 4
4. Parameter calibration ... 4
5. Modelling protein synthesis, compared to literature data ... 5
6. References ... 6


#### 1. Updating the circadian clock, starch, and photoperiod response models

##### 1.1 Photoperiod response model

The circadian clock controls the timing of flowering by regulating the expression of the *FT* gene through the photoperiod pathway. The photoperiod response was previously modelled in the Arabidopsis Framework Model version 1 (FMv1) (Chew *et al.* 2014) by including the model from Salazar et al 2009 (Salazar *et al.* 2009). However, this model includes an older circadian clock model (Locke *et al.* 2005) that does not explicitly represent the relevant clock components *PRR9* and *PRR7*. We therefore replaced the Salazar model with our most recent, Seaton-Smith model of the photoperiod pathway (Seaton *et al.* 2015). This brings several advantages. First, the Seaton-Smith model includes additional understanding of the photoperiod response mechanism, such as the regulation of CO protein stability by FKF1 (Song *et al.* 2012). Second, it is based upon the same circadian clock model (Pokhilko *et al.* 2012) as the clock-starch model that we introduce in Section 1.2, below. Third, the clock model includes *PRR9* and *PRR7*, allowing explicit simulation of the *prr9prr7* mutation (see section 1.2.2). Fourth, the Seaton-Smith model represents circadian regulation of hypocotyl elongation via the *PIF* transcription factors, allowing the FMv2 to represent this canonical clock phenotype.

As in the Salazar and FMv1 models, the photoperiod response model in the FMv2 interacts with the phenology model through the control of *FT* transcript expression. The important characteristic is '*FTarea*', the integrated *FT* level over the course of a 24h day. *FTarea* controls the *Photoperiod* component of the phenology model through the expression:

$$Photoperiod = a + b\left[\frac{c^n}{c^n + FTarea^n}\right] \qquad (1)$$

In order to utilise this connection with the new circadian clock model, the parameters *b, c* and *n* were chosen so that this function matched the original photoperiod function given by Chew et al 2012 (Chew *et al.* 2012), as was done previously for the connection from the older clock model in the FMv1 (Chew *et al.* 2014).

#### 1.2 Circadian control of starch turnover

The circadian clock controls the rate of starch degradation during the night in light:dark cycles (Graf *et al.* 2010, Scialdone *et al.* 2013). The molecular mechanisms responsible for this control have not been identified, but our recent work identified simple, plausible mechanisms (Seaton *et al.* 2014). These were formalised in mathematical models that were evaluated by comparison to a wide range of experimental data (e.g. the change in starch turnover when dusk arrives ~4 hours early). In Seaton et al 2014 (Seaton *et al.* 2014), three models were described in detail, named Model Variants 1, 2 and 3. Of these, Model Variants 2 and 3 provided the best match to experimental data, while Model Variant 1 was shown to have several limitations. Since Model Variants 2 and 3 provided quantitatively similar predictions over a range of conditions, and Model Variant 2 is simpler (6 fewer parameters and 2 fewer regulatory links from the circadian clock), we chose to integrate Model 2 with the FMv2.

##### 1.2.1 Starch model structure

In order to incorporate this control of starch turnover with the FM, we treat the starch component *S* as a measure of starch concentration (rather than absolute quantity per plant). Thus, this is taken as:

$$S(t) = r\frac{C_{starch}(t)}{C_{shoot}(t)} \qquad (2)$$

Where $S$(t) is starch concentration variable used in the model of starch turnover, $C_{starch}$(t) and $C_{shoot}$(t) are the carbon in starch and in the shoot biomass respectively, and $r$ is a scaling factor used to bring $S$(t) to a similar range of concentrations to those used in the original model construction (Seaton *et al.* 2014). Note, the control of starch synthesis by the species *Y* is disregarded, as starch synthesis is modelled as a photoperiod-dependent fraction of photoassimilate (see Section 2).

This model runs in hourly timesteps throughout the day and night, but controls starch turnover only during the night. The discrete timestep remains as in the original Carbon Dynamic Model and in FMv1, with the implicit assumption that effects within a timestep are negligible. The starch concentration (i.e. $S(t)$) is calculated at the start of the timestep, and the change in starch levels by the end of the hour is then given by:

$$\Delta C_{starch}(t) = \frac{C_{shoot}(t)\Delta S(t)}{r} \qquad (3)$$

Where $\Delta S(t)$ denotes the change in starch concentration across the hour of the simulation timestep. Total starch carbon at the following timepoint is then updated according to:

$$C_{starch}(t+1) = C_{starch}(t) + \Delta C_{starch}(t) \qquad (4)$$

$\Delta C_{shoot}$(t) is calculated subsequently within each timestep, based upon the remaining sugar level after respiration and allocation to the roots (Supporting Information Figure 2).

##### 1.2.2 Simulating *lsf1* and *prr7prr9* mutant genotypes

In order to simulate the circadian clock mutant *prr7prr9*, we set to 0 the clock parameters $q_3$, $n_4$, $n_7$, $n_8$, and $n_9$, which control the multiple aspects of the transcription rate of *PRR7* and *PRR9*. Model simulations predicted ~70% turnover of starch in the mutant, in agreement with experimental data (Fig. 3a and Supporting Information Figure 1d).

All other parameter values were calibrated as described in Section 4, below (Supporting Information Fig.4), and are shown in Supporting Information Table 1. The starch degradation rate parameter in FMv1 (*sta_turnover*) is not required in FMv2, where the starch degradation rate is computed by the clock-regulated starch model. In order to simulate the *lsf1* mutant, the parameters $k_{d,S}$ and $k_{d,T,2}$ in this model were set to 10 and 0.018, respectively, calibrating simulated starch to our experimental data. This allowed the model to match the experimentally observed starch turnover in our experiment 1 (Fig. 3g) and in literature data (Comparot-Moss *et al.* 2010) (Supporting Information Figure 1c). Where *prr7prr9* showed a mild starch phenotype in experiment 2, *sta_turnover* was calibrated as described in Supporting Information Fig. 4; the same model was used to compare all genotypes in the experiment.

#### 2. Revision of Starch synthesis

In the original Carbon Dynamic Model (CDM) (Chew *et al.* 2014, Rasse and Tocquin 2006), starch is synthesised at a rate that is the sum of a baseline rate and an 'overflow' rate. The baseline rate is a fixed proportion of the photoassimilate. The rest of the photoassimilate is first converted into soluble sugars which are used for growth and respiration. As growth demand is limited to a maximum value, any excess photoassimilate is converted into starch, through the 'overflow' rate.

Our previous work (Chew *et al.* 2014, Sulpice *et al.* 2014) showed that the 'overflow' mechanism is not always applicable, especially when plants are grown in short-day conditions (Figure 1e). Results suggested that starch is synthesised at a photoperiod-dependent fixed rate that is much higher than the baseline, and any excess photoassimilate remains as sugars. This ensures that plants store sufficient starch to last the night. We therefore re-routed the carbon flow based on this finding.

To determine the photoperiod-dependent starch synthesis rate, we first calculated the fraction of measured net assimilate partitioned to starch using our previous data(Sulpice *et al.* 2014) and the equation below:

$$F_S = \frac{S_{ED} - S_{EN}}{A_N \times P} \qquad (5)$$

where

| | | |
|---|---|---|
| $F_S$ | = | Fraction partitioned to starch |
| $S_{ED}$ | = | Starch level at ED |
| $S_{EN}$ | = | Starch level at EN |
| $A_N$ | = | Net assimilation rate per hour |
| $P$ | = | Photoperiod |

It has been reported that under low light conditions, most of the flux control through the pathway of starch synthesis resides in the reaction catalysed by AGPase (Neuhaus and Stitt 1990). Since most lab experiments are conducted under low light, we therefore also tested the relation between the fraction partitioned to starch and AGPase activity. If the total amount of starch accumulated over the light period is proportional to daily AGPase activity (averaged between ED and EN), the fraction is given by:

$$F_S(P) = \frac{k[AGPase_{average}(P)]}{A_N \times P} \quad (6)$$

where $k$ is the proportional constant. We determined the value of $k$ using data from 12-hr photoperiod as the reference. We found a strong linear relation between the fraction of measured net assimilate and photoperiod (Supporting Information Figure 2). This relation is therefore used in the FMv2 to determine starch synthesis rate, *StaSyn*, as follows:

$$StaSyn = A_N \times (-0.0296P + 0.7157) \quad (7)$$

#### 3. Addition of carbon pool for malate and fumarate

Malate and fumarate can be interconverted in the tricarboxylic acid cycle, so they are considered together in a single pool. The dynamics of this pool is modelled in a manner similar to starch except for the regulation of degradation rate by the clock. In the daytime, a fixed proportion of the photoassimilate is converted to starch, malate and fumarate, while sugar level is allow to fluctuate depending on the carbon excess. At night, malate and fumarate are consumed with a linear rate, while starch degradation rate is controlled by the clock sub-model (see Supporting Information Figure 2 and Section 1.2). For simplicity, we model a direct conversion of carbon from malate and fumarate into sugar at night, omitting the intermediate metabolic reactions.

#### 4. Parameter calibration

Results in our previous studies (Chew *et al.* 2014, Sulpice *et al.* 2014) suggested that carbon dynamics in plants are flexible and plants adjust processes like photosynthesis, starch synthesis and starch degradation rate depending on the environment. The aims of our study were to test if the dynamics of the different carbon pools can be quantitatively balanced over the timescale of vegetative growth, and how genetic regulation that modifies these dynamics affects plant growth. It is therefore necessary that the model first matches quantitatively the carbon pool data for wild-type plants for each study, reflecting the shared, environmental effects experienced by all genotypes. Model simulations and data can then be compared to test whether the model correctly predicts the growth of the wild-type plants, and to identify genetic effects on both metabolites and growth in the mutants.

To achieve this, we used parameter values measured or calculated from our data wherever possible. Mutants were then simulated by eliminating *PRR7* and *PRR9* gene expression in the clock submodel, and using genotype-specific parameters for processes that the clock in the FMv2 does not control (the organic acid pool and water content). We calibrated the following parameters to the data measured in the Col-0 control plants for each study (workflow illustrated in Supplementary Figure 4; parameter values in Supporting Information Table 1):

- photosynthesis rate was adjusted by introducing an efficiency factor relative to the default (values 0.88 and 0.90 in experiments 1 and 3, 0.79 in experiment 2)
- starch synthesis rate was adjusted by introducing an efficiency factor relative to the default (values 0.51 – 0.60)

We tuned starch synthesis and photosynthesis rates for each experiment to match the measured end-of-day level in the Columbia wild-type and used the same values in all genotypes (Supporting Information Table 1). Starch turnover was simulated by the clock-controlled starch submodel (Section 1.2), which reproduced experimental measurement of percentage turnover in experiments 1 and 3. In the altered temperature of experiment 2, starch degradation was greatly reduced, so we used the linear starch degradation introduced in the FMv1 (Chew *et al.* 2014) to reproduce starch turnover to the observed, end-of-night level.

We next calibrated the parameter values for the new carbon pool that represents malate and fumarate (MF) using Col data as follows:

- The initial level of this pool was set as 0.4 of initial starch level, based on the ratio measured in the literature (Chia *et al.* 2000)
- MF synthesis was set as a fixed fraction of starch synthesis (value 0.2 in all experiments, all genotypes)
- MF turnover was set as the fraction of dusk level consumed (values 0.6 or 0.7 in Col and *lsf1*; 0.25 or 0.21 in *prr7prr9*).

In each experiment, we did not find genotypic differences in photosynthesis when expressed per unit area, but there was a tendency toward increased photosynthesis in *prr7prr9* when expressed per gram fresh weight (Supplementary Figure 5). Even though we used the same photosynthesis efficiency for all genotypes, we found that the model could reproduce this increased photosynthetic rate per gram fresh weight in the mutant plants, due to the lower water content measured in *prr7prr9* (Supplementary Figure 9k). This reinforced the importance of including water content as a genotype-specific parameter in our model, since metabolites are measured per unit fresh weight.

As expected, we found variation in photosynthesis efficiency between experiments. In particular, the photosynthesis per unit area was higher for all genotypes in Experiment 2. As a result, the model underestimated these, but reproduced the values when expressed per unit fresh weight, suggesting a difference in the specific leaf area in this experiment that was consistent with the measured values (Supplementary Figure 9j).

The gibberellin experiment was not simulated, because the required calibration data were not all available.

#### 5. Modelling protein synthesis, compared to literature data

The biomass prediction in the FMv2 implies minimal budgets for the nutrient constituents of biomass, which are effectively predictions that can be compared to published experimental data. For example, $^{13}CO_2$ labelling has allowed quantification of the relative rates of protein synthesis in the light and dark during light:dark cycles (Pal *et al.* 2013), and of rates of protein turnover (Ishihara *et al.* 2015).

The model does not include protein as a distinct component of the synthesised biomass. However, since the protein fraction of biomass is relatively constant across the course of a day (for example, see Pyl et al 2012 (Pyl *et al.* 2012)), and protein turnover has been measured, it is possible to calculate an implied rate of protein synthesis for a given model simulation (as done experimentally in Ishihara et al 2015 (Ishihara *et al.* 2015)). In particular:

$$ProtSyn(t) = (Gr(t) + Turn)\ Prot \qquad (8)$$

where *ProtSyn*(*t*) is the calculated rate of protein synthesis at time *t*, in units of gProtein $gFW^{-1}\ h^{-1}$. *Gr*(*t*) is the relative growth rate (= *(Biomass(t)-Biomass(t-1))/Biomass(t)*), in units of hr$^{-}$

[1]. *Turn* is the rate of protein turnover, measured as 0.0014 $hr^{-1}$ (average of measurements by Pulse-Chase labelling (Ishihara *et al.* 2015)). *Prot* is the protein content, measured as 0.0169 gProtein $gFW^{-1}$ in (Ishihara *et al.* 2015).

Simulating the conditions used in (Ishihara *et al.* 2015) for wild-type plants shows that carbon biomass growth rates in the model predict a 3.3-fold increase in the rate of protein synthesis during the day, compared to during the night. This is in excellent agreement with experimental data which showed a 3.1-fold increase (Ishihara *et al.* 2015, Pyl *et al.* 2012).
